# Perturbation of transmembrane 6 superfamily member 2 expression alters lipid metabolism in a human liver cell line

**DOI:** 10.1101/2021.08.04.455062

**Authors:** Asmita Pant, Yue Chen, Annapurna Kuppa, Xiaomeng Du, Brian D Halligan, Elizabeth K Speliotes

## Abstract

**Background:** Nonalcoholic fatty liver disease (NAFLD) is caused by accumulation of excess lipids in hepatocytes. Genome wide association studies have identified strong association of NAFLD with non-synonymous E167K amino acid mutation in transmembrane 6 superfamily member 2 (TM6SF2) protein. The E167K mutation affects TM6SF2 stability and its carriers display increased hepatic lipids levels and lower serum triglycerides. While similar phenotype is evident in mice with TM6SF2 knockdown, effects of TM6SF2 on hepatic lipid metabolism is not completely understood.

**Methods:** Here, we overexpressed wild-type or E167K variant of TM6SF2 or knocked down *TM6SF2* expression in lipid-treated Huh-7 cells and used biochemical assays, untargeted lipidomic analysis, RNAseq transcriptome analysis and high-throughput fluorescent imaging to determine changes in lipid metabolism.

**Results:** Both knockdown and E167K overexpression increased acylglyceride levels which was decreased by wild-type TM6SF2 overexpression. Further, mean intensity of individual lipid droplets was increased by E167K overexpression and knockdown while wild-type TM6SF2 had no effects. We also observed lipid chain remodeling for acylglycerides by TM6SF2 knockdown leading to a relative increase in species with shorter and more saturated side chains. RNA sequencing revealed differential expression of several lipid metabolizing genes, including genes belonging to AKR1 family and lipases, primarily in cells with TM6SF2 knockdown.

**Conclusion:** Taken together, our data shows that overexpression of TM6SF2 gene or its loss-of-function changes hepatic lipid species composition and expression of lipid metabolizing genes. Further, overexpression of E167K variant and TM6SF2 knockdown similarly increased hepatic lipid accumulation and lipid droplets profile further confirming a loss-of-function effect for variant.

## Introduction

Non-alcoholic fatty liver disease (NAFLD) is one of the most common chronic liver diseases in the world^1^. NAFLD pathology is variable and can range from simple steatosis to steatohepatitis and cirrhosis^2^. No effective treatments exist for NAFLD making it a large unmet medical need. A better understanding of causes of NAFLD is important for developing novel therapeutic strategies for this disease. NAFLD is heritable and polygenic as evident by a growing list of NAFLD-related single nucleotide polymorphic risk variants identified from genome-wide association studies (GWAS)^3^. One NAFLD associated variant rs58542926, falls in the transmembrane 6 superfamily member 2 (TM6SF2) gene and causes a nonsynonymous glutamic acid to lysine substitution at the amino acid residue 167 (E167K)^4^. Presence of the E167K variant in humans is associated with higher hepatic lipid content and has been shown to be a strong genetic risk factor for NAFLD, fibrosis and cirrhosis^4–8^.

*In vitro* studies suggest that TM6SF2 E167K results in significantly reduced expression of TM6SF2 protein in liver^4,9^. Knocking out TM6SF2 in mice or inhibition of TM6SF2 gene expression in hepatocytes recapitulates human phenotype of E167K carriers suggesting that the TM6SF2 E167K variant acts by a loss-of-function mechanism^4,10^. The gene is predominantly expressed in the liver and small intestines and is mainly localized to the endoplasmic reticulum (ER) and the ER-Golgi intermediate compartment, where triglyceride (TG) rich lipoproteins (TRLs) are assembled^4,10^. TM6SF2 may have a role in the assembly of very-low density lipoproteins (VLDL) and studies have reported impaired lipidation of the VLDL particles and lower levels of plasma VLDL-TGs in human E167K carriers and in mice with TM6SF2 knockout^11–15^. The mechanism by which E167K causes hepatic steatosis, steatohepatitis, and fibrosis is less understood. While increase in TGs is one of the main hepatic phenotypes associated with TM6SF2 variant, how increased or decreased function of *TM6SF2* affects lipid droplet properties, expression of lipid metabolizing genes or cellular lipid composition is not completely understood. Indeed, many argue that a neutral lipid like triglyceride is likely not the cause of NAFLD-related liver damage and such damage may be caused by the accumulation of more hepatotoxic lipids such as saturated fatty acids, diacylglycerols, ceramides, lysophophatidyl choline, phosphoinositol, and free cholesterols which may also accumulate in NAFLD albeit at lower amounts than TG^16,17^.

Here, we over expressed wild-type or E167K variant forms of TM6SF2 or knocked down TM6SF2 in lipid-treated human hepatoma Huh-7 cells. We used biochemical assays, untargeted lipidomic analysis, RNAseq transcriptome analysis and high-throughput fluorescent microscopic image analysis to determine the cellular changes associated with altering TM6SF2.

## Results

### Overexpression of TM6SF2 (E167K) and knockdown of TM6SF2 increases acylglycerides in Huh-7

We characterized the cellular changes resulting from the overexpression of wild-type or E167K variant or knockdown of TM6SF2 in Huh-7 cells. We generated Huh7-cell lines with stable overexpression of either the wild-type TM6SF2 or its E167K variant through a strong pCMV6 promoter. We also established a stable TM6SF2 knockdown cell lines using TRC pLKO1.0 shRNA lentiviral clones. As controls for knockdown and overexpression experiments, we also constructed cell lines with an integrated lentivirus expressing a non-targeting shRNA and or an empty pCMV6-Entry vector. TM6SF2 gene expression was increased ~ 10-fold and 8-fold for wild-type or E167K variant overexpression respectively (Figure 1A) whereas TM6SF2 mRNA levels were reduced by 75% with TM6SF2 knockdown (Figure 1B). Knockdown and overexpression efficacy was verified using Western blot and was congruent with gene expression (Figure 1C). Experimental and control cells were lipid starved and then were treated with oleic acid for 24 hours before measuring intracellular acylglycerides (TG, DG and MG).

**Figure 1:**
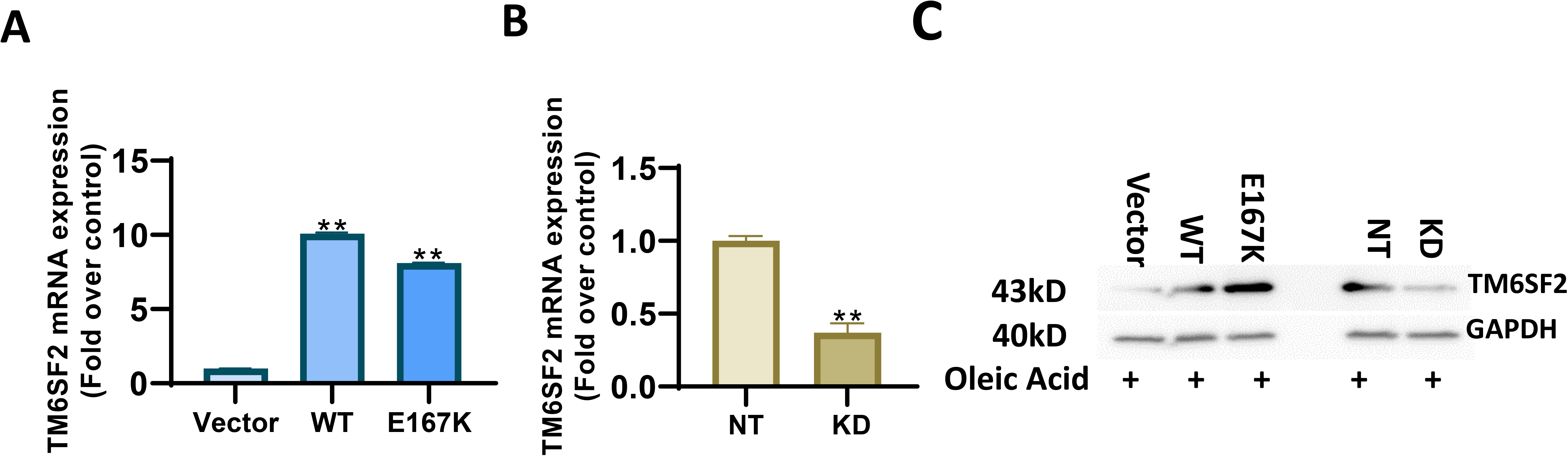
Validation of *TM6SF2* gene knockdown and TM6SF2 overexpression in Huh-7 cells. Quantitation of TM6SF2 mRNA expression using RT-qPCR in (A) overexpression and (B) knockdown Huh-7 cell lines treated with 200 μM oleic acid for 24 hours. Data presented as mean+ S.D. from 3 independent experiments, ** significantly different than control (vector or NT), *P* < 0.01. C) Western blot analysis of TM6SF2 protein levels in overexpression and knockdown Huh-7 cell lines treated with 200 μM oleic acid for 24 hours. Wild-type TM6SF2 (WT), E167K mutant variant TM6SF2 (E167K), pCMV empty vector control (Vector), non-targeted shRNA control (NT), and TM6SF2 knockdown (KD).

Overexpression of the E167K variant as well as knockdown of TM6SF2 both increased the total intracellular acylglyceride levels by ~1.8-fold whereas overexpression of wild-type lowered the acylglyceride levels by 52 % (Figure 2A, E).

**Figure 2:**
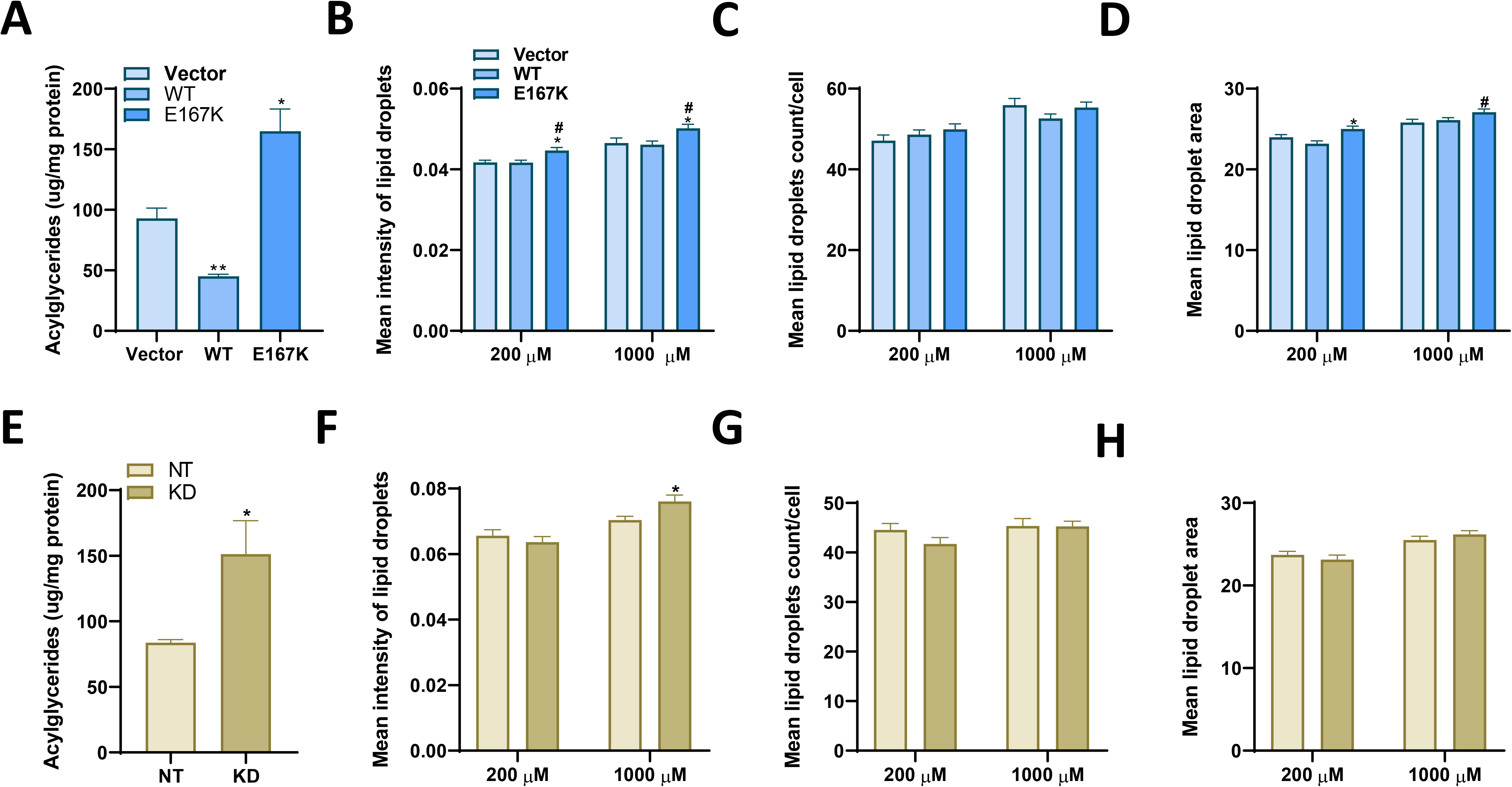
Effects of *TM6SF2* genotype on intracellular lipid accumulation and lipid droplet profile in oleic acid treated Huh-7 cells. (A, B) Biochemical measurement of intracellular acylglyceride concentrations (TG, DG, MG) in Huh-7 cell lines treated with 200 μM oleic acid for 24 hours. Quantitation of (C, D) mean intensity of lipid droplets, (E, F) mean area of lipid droplets (G, H) number of lipid droplets per cell in Huh-7 cell lines treated by 200 μM or 1000 μM oleic acid for 24 hours and stained with Hoechst (nuclei) and LipidTox Green (neutral lipids). (A-G) Data presented as mean+ S.E.M. from 3 independent experiments, * significantly different than control cells (Vector or NT), \**P* < 0.05, ** *P* < 0.01,^#^ significantly different than wild-type TM6SF2 overexpression cells (WT),^#^ *P* < 0.05. Wild-type TM6SF2 (WT), E167K mutant variant TM6SF2 (E167K), pCMV empty vector control (Vector), non-targeted shRNA control (NT), and TM6SF2 knockdown (KD).

### Overexpression of TM6SF2 E167K and knockdown of TM6SF2 increases lipid droplet intensity in Huh-7

Next, we measured the mean intensity and area of individual lipid droplets and the number of lipid droplets per cell in Huh-7 cells overexpressing wild-typeTM6SF2, TM6SF2 (E167K) or in cells with TM6SF2 knocked down along with their respective control cell lines. Mean intensity of the lipid droplets in cells with E167K variant overexpression was significantly higher at both 200 or 1000 μM oleic acid concentrations (Figure 2B) and in TM6SF2 knockdown cells at 1000 μM oleic acid concentrations (Figure 2F) compared to their respective controls at the same concentrations of oleic acid. Similarly, the mean area of lipid droplets was significantly increased by E167K variant overexpression at 200 and 1000 μM oleic acid concentrations (Figure 2D) while neither wild-type overexpression or knockdown of TM6SF2 had any effect on mean lipid droplet area (Figure 2D, H). We did not observe any significant change for lipid droplet number per cells in our overexpression or knockdown cells when compared to their respective control cells across all treatment conditions (Figure 2C, G). Compared to wild-type overexpression, E167K overexpression resulted in lipid droplets with significantly higher mean intensity at both oleic acid concentrations (Figure 2B) and mean droplet area at 1000 μM (Figure 2D) while number of lipid droplets per cell was not significantly different (Figure 2C). Taken together, results from our biochemical and image assays suggests that TM6SF2 knockdown and E167K overexpression produces similar effects on acylglyceride accumulation and lipid droplets in oleic acid treated Huh-7 cells. Since the E167K variant is considered to be a loss of function mutation, its overexpression may cause unexpected effects. For further experiments we focused on characterizing the effects of loss or overexpression of TM6SF2 function.

### Differential effects of TM6SF2 genotypes on hepatic lipid profile

Although accumulation of TGs is the major phenotype in E167K carriers or when TM6SF2 is knocked down, neutral lipids like TGs may not cause NAFLD-associated liver damage, but rather the accumulation of other more toxic lipids may determine severity and progression of NAFLD. To determine what lipid species were altered when TM6SF2 was perturbed we carried out untargeted lipidomic analysis by liquid chromatography coupled to high-resolution mass spectrometry (LC-MS/MS). Approximately 350 different lipid species distributed across 17 classes were characterized. Unsupervised hierarchical clustering of lipid species across all samples showed that replicate experimental and control samples clustered as groups, showing good concordance between replicate runs (Supplementary Figure 1). Both knockdown of TM6SF2 and wild-type overexpression significantly (adjusted *p* value < 0.05 and log fold change > 1 or < −1), changed the abundance of 30 and 27 lipid species respectively when compared to their control cells (Supplementary table 1). Most of the changes were observed across lipids classes of TGs, DGs, phospholipids (PC, PE, PS, and PG), sphingomyelins, lyso phospholipids and plasmenyl phospholipids and the detail changes are shown in Supplementary Table 1.

Wild-type overexpression significantly changed the abundance of 27 lipid species when compared to the vector control (Supplementary table and Figure 3A). The two lipid species with the largest log_2_ fold change were plasmenylPC 44:4, which increased approximately by 11 fold with wild-type overexpression and plasmenylPE 34:0, which decreased by 3.4 fold. The two lipid species with the smallest adjusted *p* value were PE 31:1 and PE 32:2 (adjusted p value of 0.002), which both increased by 2.8 and 3.4 fold respectively with wild-type overexpression. Two triglyceride species, TG 52:6 and TG 60:5 were both decreased by approximately 2.7 fold and three diglyceride species, DG 36:3, DG 38:2, and DG 38:4, were also decreased by approximately 2.4 fold by wild-type overexpression. In the knockdown comparison (Figure 3B), the highest log fold change was found with lysoPE 20:3, which increased by approximately 4 fold with knockdown and PC 42:9, which decreased approximately 4.6 fold with knockdown. Five lipid species with the lowest adjusted *p* value (7 × 10^−5^) were lysoPC 18:0, lysoPC 20:3, lysoPE 20:3, PG 38:7, and plasmenylPE 38:3 all of which increased in the knockdown by 2 to 3 fold. Four other lipid species with the lowest adjusted *p* value (7 × 10^−5^) were SM 33:0, TG 58:8, TG 60:7, and TG 62:7 all of which decreased in the knockdown by approximately 2.1 fold. There were 7 triglyceride species that were significantly changed between the knockdown and non-targeted control cells. In the knockdown, TG 52:1 and TG 58:1, were increased by approximately 2 fold and TG 56:1, TG 58:7, TG 58:8, TG 60:7, and TG 62:7 were decreased by approximately 2 fold. The two diglycerides, DG 31:0 and DG 34:0, were increased by 2.1 and 2.3 fold in the knockdown

**Figure 3:**
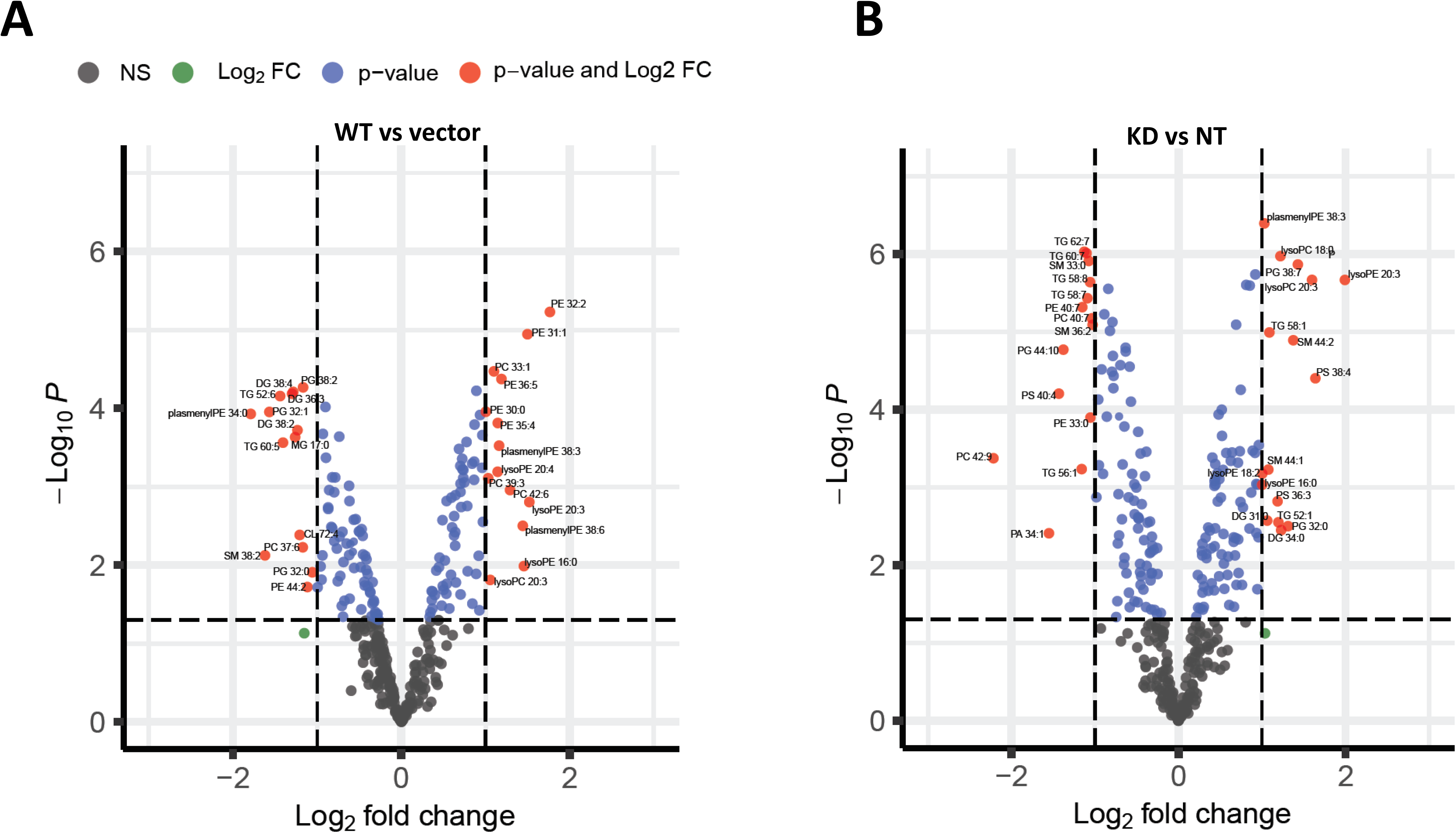
Effects of TM6SF2 genotypes on hepatic lipid profile. Volcano plots showing relative abundance of lipid species identified through untargeted lipidomic analysis. X-axis is the log_2_ fold change of lipid species abundance in Huh-7 cells for (A) wild-typeoverexpression compared to vector control (B) TM6SF2 knockdown compared to non-targeting control. For both A and B, Y-axis represents the adjusted *p* value. Negative values indicate downregulated lipids while positive values upregulated lipids in the overexpression or knockdown cells compared to their respective controls. Vertical dotted lines represent the threshold for log_2_ fold change (> 1 or < −1) and horizontal dotted line represent the threshold for adjusted *p* value (*P* < 0.05). Only the lipid species that met the threshold criteria for both p-value and log fold change are labelled (red dots). Values for log fold change and significance for lipids that meet both threshold are shown in detail in Supplementary Table 1. Wild-type TM6SF2 (WT), pCMV empty vector control (Vector), non-targeted shRNA control (NT), and TM6SF2 knockdown (KD).

We also analyzed lipidomics results for changes in side chain length and unsaturation across observed lipid classes. Heat maps showing the distribution of unsaturation and chain length for each lipid class is shown in supplemental Figure 2. Overall, there were small changes in the distribution of chain length and saturation for most lipid classes. The most significant changes were observed for DGs and TGs (Figure 4). Wild-type TM6SF2 overexpression generally decreased the abundance of TG and DG species across all saturation and chain lengths (Figure 4A, C). In contrast, TM6SF2 knockdown significantly increased abundance of more saturated TGs and DGs with shorter side-chain length while the abundance of TGs and DGs with higher side-chain lengths and unsaturation was decreased (Figure 4B, D). Since some of the highest levels of changes in abundance for lipid species were observed for lyso lipids, particularly the lysoPCs and lysoPEs in the TM6SF2 knockdown cells, we also looked at the relative changes in abundance of lyso lipids by chain length and unsaturation (Supplementary figure 2). However, we did not observe any major shift in the distribution of chain saturation or side-chain length for lyso lipids, which is in contrast to acyl lipids where it seemed that the change in abundance was correlated to a shift in chain length and unsaturation instead of increase in a specific lipid species.

**Figure 4:**
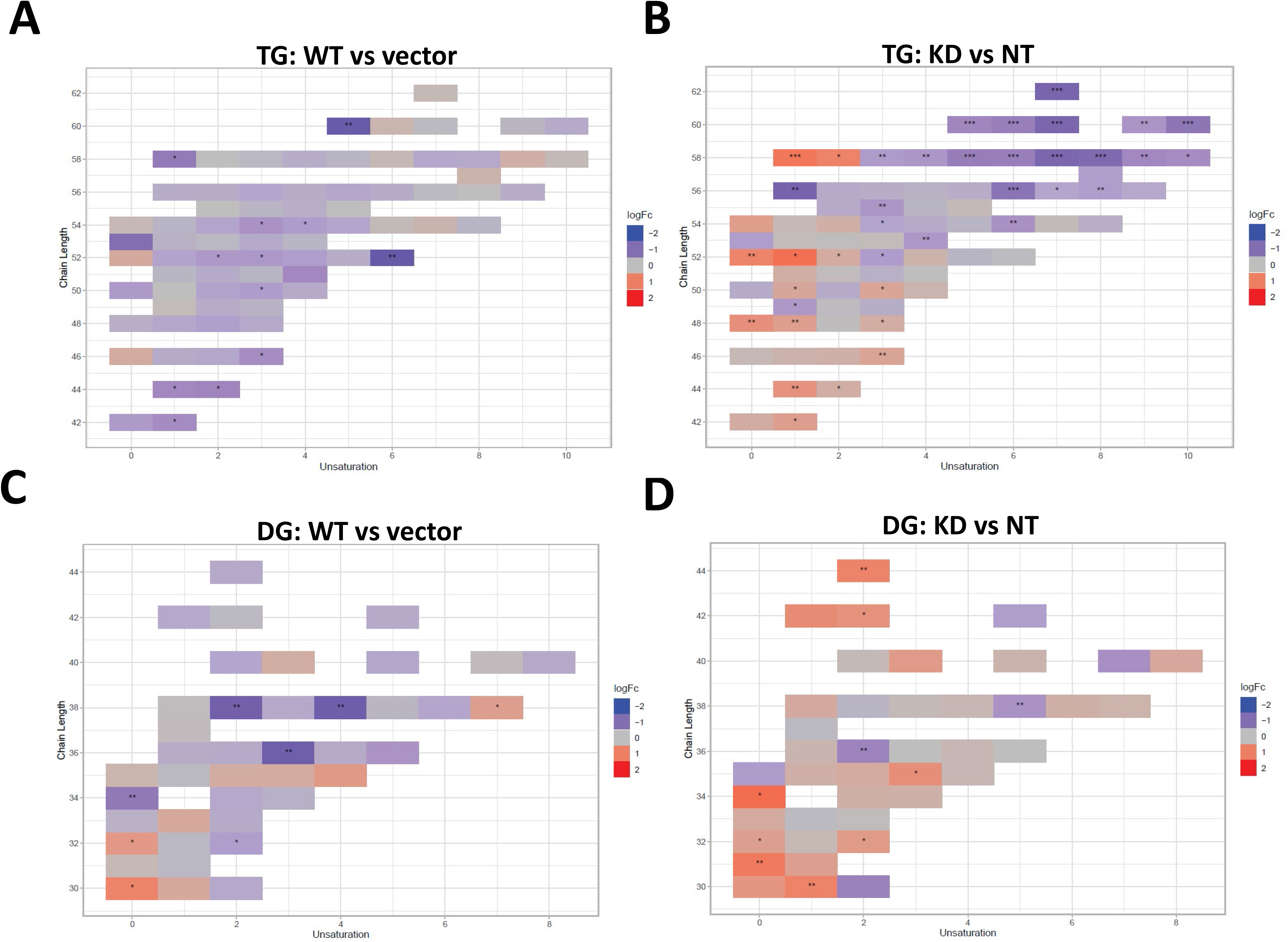
Knockdown of TM6SF2 decreases longer and more unsaturated TGs and DGs. Heatmap shows the distribution of individual TGs (A, B) and DGs (C, D) across total degree of unsaturation and the side chains lengths. Each box represents an individual lipid species where the colors indicate the relative ratio of the lipid (red: increased; white: no difference; blue: decreased) in (A, C) wild-typeoverexpression (n=3) compared to vector control (n=3) and (B, D) TM6SF2 knockdown (n=5) versus non targeting control (n=4). Lipids with statistically significant difference between the experimental and control cells are highlighted. * *P* <0.05, ** *P* <0.01, *** *P* <0.001. Wild-type TM6SF2 (WT), pCMV empty vector control (Vector), non-targeted shRNA control (NT), and TM6SF2 knockdown (KD).

Taken together, our results show that knockdown of TM6SF2 increased DGs, lysoPC, and lysoPE lipids, decreased PCs and PEs, and also caused a shift in acyl glyceride lipid side chains to become more saturated with shorter chain lengths. Wild-type TM6SF2 overexpression decreased TGs, DGs, MGs, and PGs, increased PCs, PEs, and lysoPEs, and had no effect on chain lengths of lipids.

### RNAseq analysis show changes in expression of genes involved in lipid metabolism in response to different TM6SF2 expression conditions

Next, we performed RNAseq analysis to determine changes in gene expression resulting from altered TM6SF2 expression. We focused our analysis on the combination of the set of 1008 genes for the combination of the 749 proteins identified by Reactome as being part of the overall human Metabolism of lipids pathway (R-HSA-556833) and the 864 human proteins identified in GO as being involved in the regulation of lipid metabolic process (GO:0019216). Volcano plots of differential expression of these genes are shown in Figure 5. Compared to empty vector, 16 genes were differentially expressed (adjusted *p* value of less than 0.05 and an absolute log fold change of greater than 0.5 or 1.4-fold change) in cells with wild-type overexpression (Figure 5A) while 96 lipid metabolizing genes were differentially expressed in the cells with TM6SF2 knockdown compared to the non-targeting control (Figure 5B). Complete sets of genes are listed in the Supplementary Table 2. Four members of the Aldo-keto reductase family (AKR1B1, AKR1C2, AKR1C4, AKR1D1) were increased from 2 to 6 fold by TM6SF2 knockdown whereas expression of AKR1C4, AKR1C2 and AKR1C1 were all decreased by at least 1.5-fold in wild-type overexpression cells. Expression of both LDLR (low-density lipoprotein receptor) and (LSR) lipolysis-stimulated lipoprotein receptor was increased by ~ 1.4-fold and of LRP2 (low-density lipoprotein receptor-related protein 2) was decreased by ~ 1.8-fold. We also observed changes in the expression of apolipoproteins such as APOA5 (Apolipoprotein A-V) and APOC3 that decreased by ~ 1.8 and ~2-fold respectively by knockdown. In contrast, decreased expression of APOA2 (~ 1.5-fold) and increased expression of APOD (~ 4-fold) were observed in cells with wild-type overexpression. Across the two comparisons, 16 different genes involved in the metabolism of lysophosphatidylethanolamine (LPE) were significantly changed. LIPH, a gene involved in LPE hydrolysis, was increased in both conditions and PNPLA2 was decreased in knock down of TM6SF2. TM6SF2 RNA expression was increased by 3.4 fold in the wild-type overexpression and reduced to 0.82 in the knockdown, consistent with the qPCR results (Figure 1A, B).

**Figure 5:**
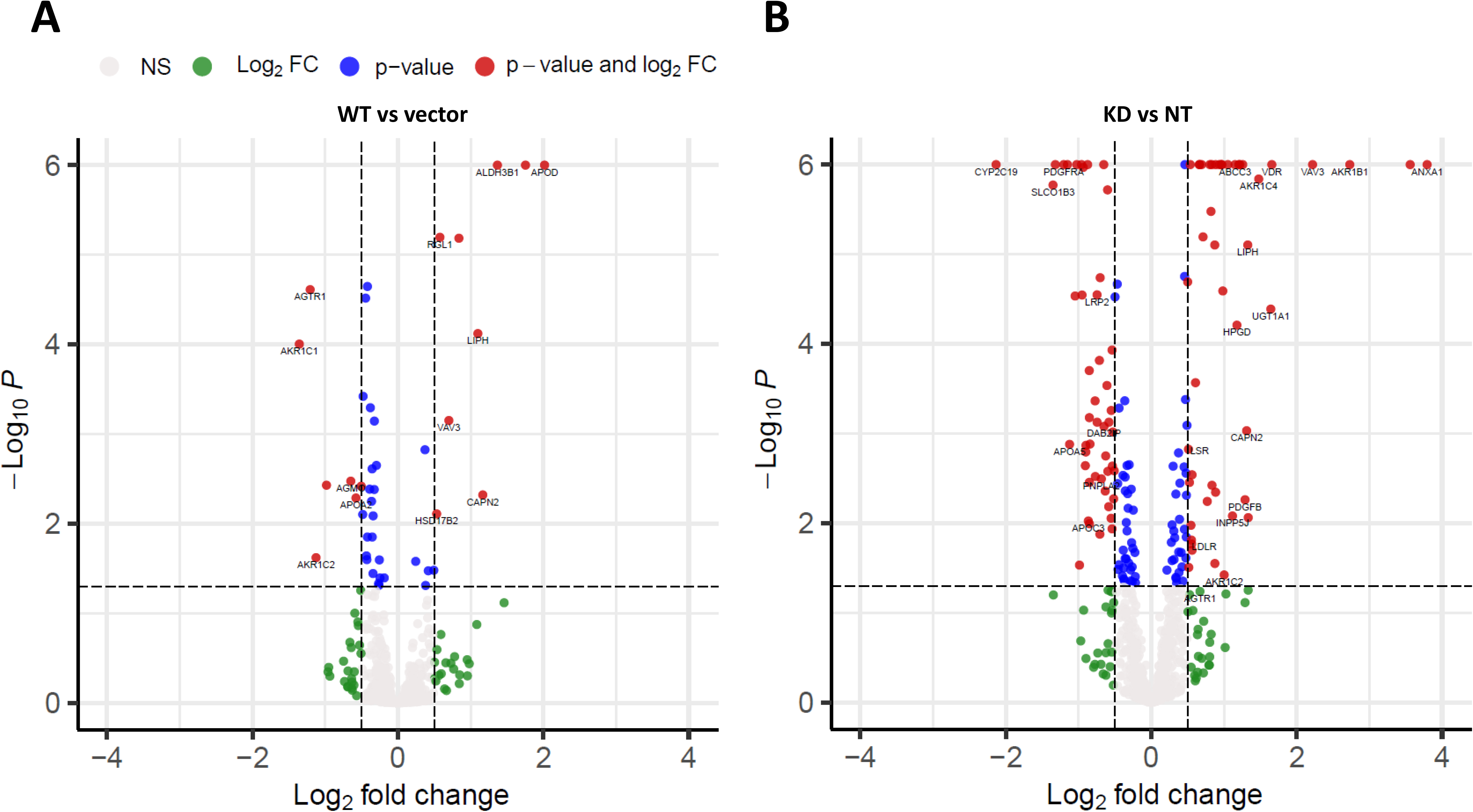
Changes in expression of lipid metabolizing genes in response to TM6SF2 overexpression or knockout. Volcano plots shows relative abundance of lipid metabolizing genes identified through RNA Sequencing. X-axis is the log_2_ fold change of gene expression in Huh-7 cells for (A) wild-typeoverexpression compared to vector control (B) TM6SF2 knockdown compared to non-targeting control. For both A and B, Y-axis represents the adjusted *p* value. Negative values indicate downregulated genes while positive values indicate upregulated genes in the overexpression or knockdown cells compared to their respective controls. Vertical dotted lines represent the threshold for log_2_ fold change (> 0.5 or < −0.5) and horizontal dotted line represent the threshold for adjusted *p* value (*P* < 0.05). Genes belonging to the lipase or the AKR1 family of genes that met the threshold criteria for both p-value and log fold change are labelled (red dots). Detailed information on the names, fold change and significance of genes meeting both threshold are shown in Supplementary Table 2. Wild-type TM6SF2 (WT), pCMV empty vector control (Vector), non-targeted shRNA control (NT), and TM6SF2 knockdown (KD).

## Discussion

The primary objective of this study was to determine the differential effects of TM6SF2 overexpression and loss on hepatic lipid metabolism, composition and lipid droplet dynamics under steatogenic conditions. We utilized four parallel experimental approaches to determine changes in total levels of acylglycerides, gene expression, individual lipid species, and lipid droplet phenotype in oleic acid-treated Huh-7 cells. Our data shows that knockdown of TM6SF2 and overexpression of the E167K mutant variant increases whereas overexpression of the wild-type lowers acylglycerol accumulation in lipid-loaded hepatocytes. While we did not see changes across lipid classes, we observed lipid chain remodeling for TGs and DGs by TM6SF2 knockdown leading to a relative increase in shorter and more saturated side chains on glycerol lipids. Importantly, TM6SF2 knockdown and overexpression lead to significant changes in the abundance of several lipid species, including phosphoplipids, lysophospholipids, and acylglycerides. RNA sequencing revealed differential expression of lipid metabolizing genes belonging to the AKR1 family and lipases with TM6SF2 knockdown causing most of the significant changes. Characterization of lipid droplets profile through high-content image analysis determined that overexpression of the E167K variant and TM6SF2 knockdown significantly increase the staining intensity and size of lipid droplets. Taken together, our data shows that overexpression of TM6SF2 gene or its loss of function significantly and differentially affects intrahepatic lipid metabolism to change lipid species composition, expression of lipid metabolizing genes and lipid droplet profile in oleic acid-treated Huh-7 cells. Further, overexpression of TM6SF2 E167K and TM6SF2 knockdown increased hepatic lipid accumulation and lipid droplets profile confirming a loss of function effect for the variant.

While data from multiple human studies have established the strong association of TM6SF2 E167K mutant variant with NAFLD, the exact mechanism how the loss of TM6SF2 function causes hepatic steatosis is not entirely clear. Study in NAFLD patients and in animal models have largely focused on understanding how the E167K variant or the TM6SF2 knockdown affects very-low density lipoprotein (VLDL) and secretion of VLDL TG, and on TGs levels in plasma or in the liver^12–15,18^. Similarly, most of the *in vitro* studies on TM6SF2 knockdown or overexpression explored the effects of gene expression modification in hepatic lipid accumulation under basal conditions and in the absence of lipid exposure^10,19^. Here we evaluated the effects of TM6SF2 overexpression or its loss of function on intrahepatic lipid metabolism under a steatogenic environment and show that increased accumulation of intracellular lipids is clearly evident in oleic acid-treated Huh-7 cells with TM6SF2 knockdown or E167K variant overexpression Additionally, in the present study, we utilized high-throughput imaging with high-content image analysis to determine changes in hepatic lipid accumulation at individual lipid droplet level. Our results show significant increases in intensity and area of individual lipid droplets with TM6SF2 knockdown or E167K overexpression suggesting presence of larger more TG rich lipid droplets in the cells. One possible mechanism behind this observation could be the previously demonstrated role of TM6SF2 on lipidation of very low density lipoproteins and data showing that its loss of function results in intrahepatic retention of TGs^13^.

TM6SF2 E167K variant has also been associated with progression of NAFLD to NASH^7,8^. Although accumulation of TGs is the major phenotype in E167K carriers, neutral lipids like TGs may not lead to NAFLD-associated liver damage, but rather the accumulation of other more toxic lipids may determine severity and progression of NAFLD. The current study, overexpression the wild-type TM6SF2 did not cause any major changes in lipid species that have been previously shown cause liver damage in NAFLD. However, TM6SF2 knockdown significantly increased in lysoPC and lysoPE, both of which were recently identified as early biomarkers of NAFLD and are increased in NALFD patients with hepatitis^20^. LysoPC has also been previously shown to be increased in liver of NASH patients^21^. However, lysoPE but not lysoPCs were also increased by wild-type overexpression, therefore, further study is needed to determine the significance of these changes.

Previous studies show that hepatic TGs or TGs extracted from the VLDLs in liver of E167K carriers have shorter and more saturated fatty acids^15,22^. In accord with these findings, we observed relative increase in the abundance of saturated acylglycerides with shorter side chain lengths in cells with TM6SF2 knockdown. In addition to TGs, previous studies also suggest that TM6SF2 knockdown may lead to a global shift in lipids in terms of saturation, including PCs, which are major membrane phospholipids^23^. At the same time, studies show variable results on TM6SF2 knockdown-mediated changes in the total abundance of hepatic PCs with data showing both decrease or increase in Huh-7 cells, no change in mice and decrease in human E167K carriers^13,22,23^. Luukkonen et. al also reported that PCs synthesis from polyunsaturated fatty acids is impaired in E167K variant carriers thereby lowering total PCs in liver. Although we did not see total class level changes for PCs in our study, abundance of two of the PC species were significantly decreased by TM6SF2 knockdown while opposite effects were observed for wild-type TM6SF2 overexpression where abundance of three out of four significantly changed PCs were increased. Similar effects were also observed for significantly changed PE species, all of which were decreased by TM6SF2 knockdown and increased by wild-type overexpression.

Members of aldo-keto reductase 1 (AKR1) family, enzymes that regulate steroid metabolism, have been previously known to play a major role in hepatocarcinoma^24^. Increasing evidence suggest that several members of the AKR1 family such as AKR1B7 and AKR1B10 might play important role in the development of NAFLD or NASH^25,26^. Here, expression of several AKR1 family of enzymes were either increased or decreased in all of our cell lines compared to controls, with most changes observed for cells with TM6SF2 knockdown. We also found differential expression of lipases, PNPLA2 and LIPH in our study along with changes in the expression of lipoprotein receptor genes (LDLR, LSR, LRP2), which could be cellular response to increased TG levels in the cells. Further investigation is needed to reveal any mechanistic association of these genes in TM6SF2-mediated changes in hepatic lipid levels.

In this study, we combined systems biology, biochemical assays and high-throughput high-content image analysis to get a more complete picture of hepatic lipid metabolism-associated changes resulting from alterations in *TM6SF2* gene expression than currently available. Our data shows some novel changes in lipid composition and expression of lipid metabolizing genes, however, the mechanism how these changes manifest and their role, if any, on hepatic steatosis and other NAFLD-related phenotypes needs further investigation. In summary, the current study shows that overexpression of wild-type TM6SF2 or loss of TM6SF2 function can effectively and differentially modulate hepatic lipid metabolism to produce significant changes in intracellular lipid levels and lipid species composition. Importantly, loss of TM6SF2 function may produce more profound and overall changes in lipid metabolic pathways under a steatogenic environment. Finally, we have shown that *in vitro* model used in this study, particularly the TM6SF2 knockdown studies, recapitulates many of the lipid-metabolism associated phenotypes that were observed in human E167K carriers. Thus, use of this *in vitro* model along with high-throughput experimental study design such as high-content image analysis used in this study could be a valuable tool to perform further functional analysis on TM6SF2 or other GWAS identified genetic modifiers of hepatic steatosis.

## Methods

### Cell line generation and experimental Design

We generated Huh-7 cell lines stably overexpressing wild-type or the E167K variant of TM6SF2 or stably knocking down TM6SF2. Detailed method for cell line generation is explained in supplementary material. Briefly, we infected Huh-7 cells with TRC pLKO1.0 shRNA lentiviral clones and selected for stable TM6SF2 knockdown cell lines using 10μg/ml puromycin. Cell lines stably overexpressing wild-type or E167K variant of TM6SF2 were produced by transformation with plasmids expressing the wild-type or variant cDNA under control of a strong CMV promoter and selection for continued antibiotic resistance using 10μg/ml blasticidin. As controls for knockdown and overexpression experiments, we also constructed cell lines with an integrated lentivirus expressing a non-targeting shRNA and or an empty pCMV6-Entry vector. For our studies, Huh-7 cells (controls or genetically modified cells) were cultured in high glucose (25mM) DMEM with 10% FBS and 1% PenStrep along in the presence of antibiotics. After 24 hours, the growth medium was replaced with DMEM containing 10% delipidated FBS. 24 hours after lipid starvation, 200μM of BSA-conjugated oleic acid was added to DMEM culture medium. Cells were harvested after 24 hours and samples were collected for RNA, protein, and intracellular triglyceride measurements and lipids were extracted for LC-MS/MS untargeted lipidomics analysis. Similarly treated cells were also fixed and stained for image analysis using high-throughput microscopy.

### Measurement of gene and protein expression

*TM6SF2* gene expression was measured by Real time quantitative RT-qPCR method, which is described in detail in the Supplementary material. Briefly, total cellular RNA was extracted from Huh-7 cells treated with oleic acid for 24 hours using TRIZOL reagent (ThermoFisher Scientific) and reverse transcribed using the Superscript VILO (Life Technologies) reverse transcriptase kit. RT-qPCR PCR was then performed using TaqMan^®^ Gene Expression Assay FAM probes for TM6SF2 gene (Life Technologies). For TM6SF2 protein estimation, cells were homogenized using the RIPA cell lysis buffer and the supernatant with total protein was collected for analysis. Protein concentrations were determined using modified Bradford Protein assay according to the manufacturer’s protocol (Thermal Scientific), size-fractionated in 12% SDS-PAGE gel, and transferred to a PVDF nitrocellulose membrane (Millipore). TM6SF2 proteins were then visualized by using the mouse polyclonal anti-TM6SF2 antibody (Abcam) and Horseradish peroxidase-conjugated goat anti-mouse IgG (Sigma-Aldrich) secondary antibody. The membrane was then stripped and levels of the GAPDH protein was measured similarly using the mouse monoclonal anti-GAPDH antibody (Proteintech) and Horseradish peroxidase-conjugated goat anti-mouse IgG (Sigma-Aldrich) secondary antibody. The blots were visualized using SuperSignal-enhanced chemiluminescence (Thermo Scientific, Rockfold, IL, USA) and chemiluminescence function on a GE Healthcare Amersham AI 600 RGB imager. Image J software (NIH) was used to analyze the intensity of TM6SF2 and GAPDH protein bands. TM6SF2 protein levels were normalized to GAPDH protein level. For detailed methods, please refer to the Supplementary materials.

### Acylglyceride assay

After harvesting, total cellular triglycerides were extracted from cells using the Folch extraction method and quantified by using a Serum Triglyceride Determination Kit (Sigma, St. Louis, MO, USA). The absorbance was read at 540nm using an Eppendorf Biospectrometer and 2.5 mg/mL glycerol was used as the standard (Sigma). The total cellular triglyceride content was normalized to total cellular protein.

### Sample preparation for untargeted lipidomics

24 hours after oleic-acid exposure, cell culture medium was removed and cells were washed twice with 1X PBS buffer. Cells were then collected by 0.05% Trypsin-EDTA lysis, counted using LOGO automated cell counter and stored at −80°C until analyzed. The lipids were extracted using a modified Bligh-Dyer method. The extraction was carried using 2:2:2 (v/v/v) water/methanol/dichloromethane at room temperature after spiking internal standards. The organic layer was collected and dried completely under the stream of nitrogen. Dried extract was re-suspended in 100 μL of 10 mM ammonium acetate.

### LC-MS/MS

Lipid extract was injected onto a 1.8 μm particle 50 × 2.1 mm id Waters Acquity HSS T3 column (Waters, Milford, MA) heated to 55°C. The column was eluted with acetonitrile / water (40:60, v/v) with 10 mM ammonium acetate as solvent A and acetonitrile / water / isopropanol (10 : 5 : 85 v/v) with 10 mM ammonium acetate as solvent B. Gradient is 0 min 40% B, 10 min 100% B, 12 min 100% B, 12.1 min 40% B, 15 min 40% B. Data were acquired in positive and negative mode using data-dependent MSMS with dynamic mass exclusion at the University of Michigan Metabolomics core. Pooled human plasma sample and pooled experimental sample (prepared by combining small aliquots of all client’s samples) were used to control the quality of sample preparation and analysis. Randomization scheme was used to distribute pooled samples within the set. Mixture of pure authentic standards is used to monitor the instrument performance on a regular basis.

### LC-MS/MS Data analysis

Lipids are identified using LIPIDBLAST package (http://fiehnlab.ucdavis.edu/projects/LipidBlast), computer-generated tandem MS library of 212,516 spectra covering 119,200 compounds representing 26 lipid classes, including phospholipids, glycerolipids, bacterial lipoglycans and plant glycolipids. Quantification of lipids is done by Multiquant software (AB-SCIEX). Data was normalized using internal standards first and cell number. Data from positive and negative ion mode runs were combined with repeats removed and filtered by RSD (<30%).

### Lipidomics Data Analysis

Combined normalized data for each run was imported into the R package lipidr (https://github.com/ahmohamed/lipidr). Data was imported as a lipidomics object, summarized, annotated, normalized using the PQN method. Lipidr was used for analysis and to generate graphs.

### High-throughput lipid droplet imaging and analysis

Huh-7 cells were plated into 96-Well optical-bottom plates (Thermo Scientific, 1256670) at the density of 5,000 cells/well in conditions similar to other studies. 24 hours after oleic acid treatment, cells were fixed in 4% paraformaldehyde for 10 minutes in RT, washed three times with 1X PBS and permeabilized with 0.1% Triton-X for 10 minutes. Intracellular lipid droplets were stained with the LipidTOX Green neutral lipid stain (1:1000; Thermo Scientific, H34475) and the nucleus was stained with DAPI (1:1000; Thermo Scientific, D1306) for 30 minutes and cells were then washed twice in PBS. Cells were imaged using the Cellomics Array Scan VT1 (Thermo Scientific) at a magnification of 20X. High-content image analysis was done using the CellProfiler software. The pipeline used for CellProfiler imaging is added as supplementary method.

## Supporting information

Supplemental cell profiler method

## Acknowledgements

We would like to sincerely thank Dr. Jonathan Sexton, Assistant Professor of Internal Medicine, Medical School, and Assistant Professor of Medicinal Chemistry, College of Pharmacy for kindly letting us access to the CellInsight CX5 High-Content Screening (HCS) Platform (Thermofisher Scientific).

