## Supplementary methods and tables for "Perturbation of transmembrane 6 superfamily member 2 expression alters lipid metabolism in a human liver cell line"

### Supplementary materials

#### Supplementary method:

**Cloning human TM6SF2 cDNA into pCMV6-Entry vector**
Human wild-type *TM6SF2* was amplified by PCR from a human cDNAs library using primers containing Mlu I and EcoR I restriction sites and was sub cloned into the pCMV6-Entry vector (OriGene, Rockville, MD) using the Gibson cloning kit with Q5 High-Fidelity DNA Polymerase (New England Biolabs, Ipswich, MA). The sequences of *TM6SF2* insert was confirmed by Sanger sequencing with 3X coverage of sequencing (UM Sequencing Core).

**Preparation of human pCMV6-TM6SF2 (E167K) mutant cDNA plasmid**
The coding sequence for the human *TM6SF2* mutant variant (E167K) cDNA was amplified from wild-type TM6SF2 cDNA plasmid (pCMV6-TM6SF2 plasmid DNA as PCR template) by PCR site-directed mutagenesis using QuikChange$^{TM}$ Site-Directed Mutagenesis Kit (Stratagene, San Diego, CA). The sequences of *TM6SF2* mutant variant (E167K) construct was confirmed by Sanger sequencing with 3X coverage of sequencing (UM Sequencing Core).

**Overexpression of TM6SF2 (wild-type and E167K mutant) genes in cultured human hepatoma cells**
Huh-7 cells were grown to 70% confluence in DMEM medium with 10% FBS plus 100 IU/ml penicillin and 100 µg/ml streptomycin. For *TM6SF2* overexpression, Huh-7 cells were transfected with the pCMV6-Entry vector expressing *TM6SF2* (wild-type or E167K mutant) genes or a control empty pCMV6-Entry vector using FuGENE transfection reagent (Thermo Scientific). Post transfection (48 hours), cells stably expressing these genes were selected with G-418 (10 µg/mL) for 72 hours. Culture DMEM medium was freshly changed and fresh G-418 was added every 2 days. After 3 days, cells stably overexpressing the *TM6SF2* (wild-type or E167K mutant) gene were collected and the total cellular RNA was extracted using TRIzol reagent (Thermal Fisher Scientific). Overexpression was measured and confirmed by RT-qPCR.

**Knockdown of endogenous TM6SF2 gene expression in cultured human hepatoma cells**
Huh-7 cells were grown to 70% confluence in DMEM medium with 10% FBS plus 100 IU/ml penicillin and 100 µg/ml streptomycin and infected with human *TM6SF2* shRNAs (clone 382425, clone 382426) lentivirus (Sigma-Aldrich Company). At 72 hours post infection, Huh-7 cells stably expressing *TM6SF2* shRNAs lentivirus vector were selected in new C-DMEM medium containing 10 µg/ml puromycin (Sigma-Aldrich Company) for 7 days. The culture medium was changed and fresh puromycin (10µg/ml) added every 2 days. After 7 days, Huh-7 cells with stable *TM6SF2* knock down were collected and total cellular RNA was extracted using TRIZOL reagent. Knockdown of *TM6SF2* mRNA was confirmed by RT-PCR.

**Real time quantitative RT-qPCR**
Total cellular RNA was prepared from stable overexpression/knockdown Huh-7 cells using TRIZOL reagent (ThermoFisher Scientific) and purified by ZYMO Clean Kit. Superscript VILO (Life Technologies) reverse transcriptase kit was used to synthesize the first strand cDNA, and reverse transcribed cDNA served as template in polymerase chain reactions. The real-time relative quantitative

PCR was performed using TaqMan® Gene Expression Assay (Life Technologies) FAM probes for TM6SF2 gene respectively with TagMan® Gene Expression Master Mix (Life Technologies) following the manufacturer's protocol. The TaqMan® ELF1 FAM Gene Expression Assay probes (Life Technologies) was used as endogenous gene controls.

**Protein extraction and Western blotting**

24 hours after oleic acid treatment, Huh-7 cells were washed twice with 1XPBS, harvested from their culture wells using 0.25% trypsin-EDTA solution (Sigma), washed with phosphate-buffered saline, and then homogenized in 0.35 ml of RIPA cell lysis buffer (3 mM 2-mercaptoethanol; 0.1 mM phenylmethylsulfonyl fluoride; 20 mM Tris, pH 7.4; 160 mM NaCl; 0.03% Tween-20; and 0.1 mM EDTA) (Fisher Scientific Company). The homogenate was then centrifuged at 10,000 x g for 30 min, and the supernatant was collected for analysis. Protein concentrations were determined using modified Bradford Protein assay according to the manufacturer's protocol (Thermal Scientific). A proportion (3X volume) of cell lysate was added to 1X volume 4X sample loading buffer to a final concentration of 1X. After heating to 100 ºC for 10 min, the proteins were size-fractionated by 12% SDS-PAGE at 100 V and then transferred to a PVDF nitrocellulose membrane (Millipore) at 35 V for overnight at 4 ºC. The membranes were incubated in TBST buffer with 3% goat serum (Thermal Fisher) at room temperature for 1 h before adding the primary antibody. Mouse polyclonal anti-TM6SF2 antibody (Abcam) was diluted (1:2500) in TBST buffer with 1.5 % goat serum and incubated with membranes for overnight at 4° C. Membrane was washed three times for 10 min each in TBST buffer. Horseradish peroxidase-conjugated goat anti-mouse IgG (Sigma-Aldrich) were diluted (1:5000) in TBST buffer plus 1.5 % goat serum and incubated with membranes for 1 h at room temperature. Membrane was subject to four 5-min washes in TBST and visualized using SuperSignal-enhanced chemiluminescence (Thermal Scientific). To strip the TM6SF2 antibody, membranes were washed with 1XPBS twice and incubated with Restore Plus Western Blot Stripping Buffer (Thermal Fisher) for 15 minutes. After washing with 1XPBS buffer one more time, membranes were blocked in TBST buffer with 3% goat serum (Thermal Fisher) at room temperature for 1 h. Then mouse monoclonal anti-GAPDH antibody (Proteintech) was diluted (1:2000) in TBST buffer with 1.5% goat serum and incubated with membranes for 1 h. Membrane was washed three times for 10 min each in TBST buffer and Horseradish peroxidase-conjugated goat anti-mouse IgG (Sigma-Aldrich) were diluted (1:5000) in TBST buffer plus 1.5 % goat serum and incubated with membranes for 1 h at room temperature. Membrane was subject to four 5-min washes in TBST and visualized using SuperSignal-enhanced chemiluminescence (Catalog No. 34580, Thermo Scientific, Rockfold, IL, USA). The blot was visualized using chemiluminescence function on a GE Healthcare Amersham AI 600 RGB imager. Image J software (NIH) was used to analyze the intensity of TM6SF2 and GAPDH protein band in the Western blot.  TM6SF2 protein levels were normalized to GAPDH protein level.

**Supplementary Figure 1**: Unsupervised Hierarchical clustering analysis for 350 lipid species.

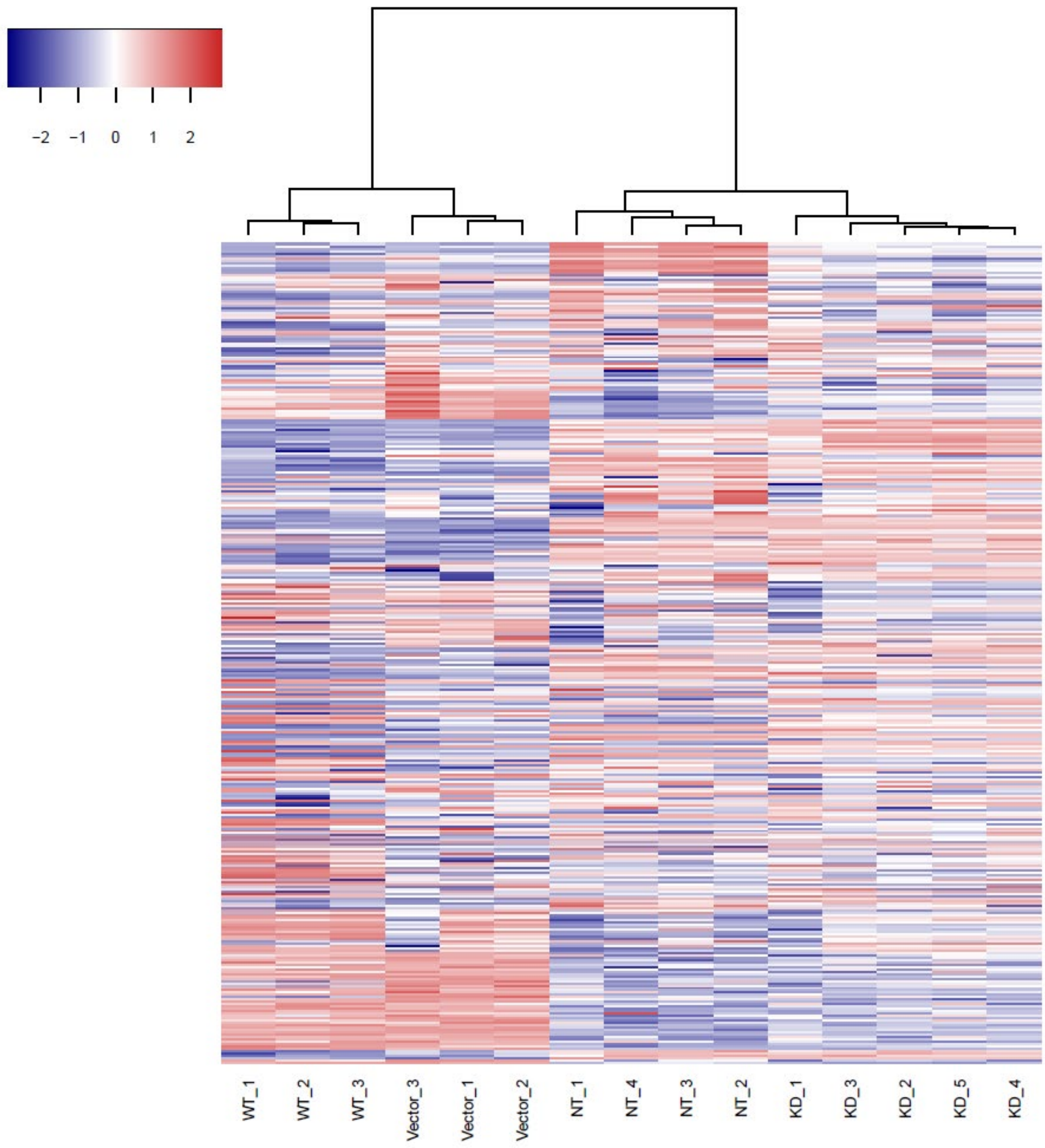


**Supplementary Figure 1:** Red color represents the abundance of each lipid species (red: higher, blue: lower). WT (wild-type TM6SF2 overexpression, Vector (empty vector control), NT (Non-targeted shRNA control), and KD (TM6SF2 overexpression). Numbers at the ends refers to each replicates.

**Supplementary Figure 2:** Heatmap of side chain length and degree of unsaturation for lipid species identified in the lipidomic analysis.

**A**

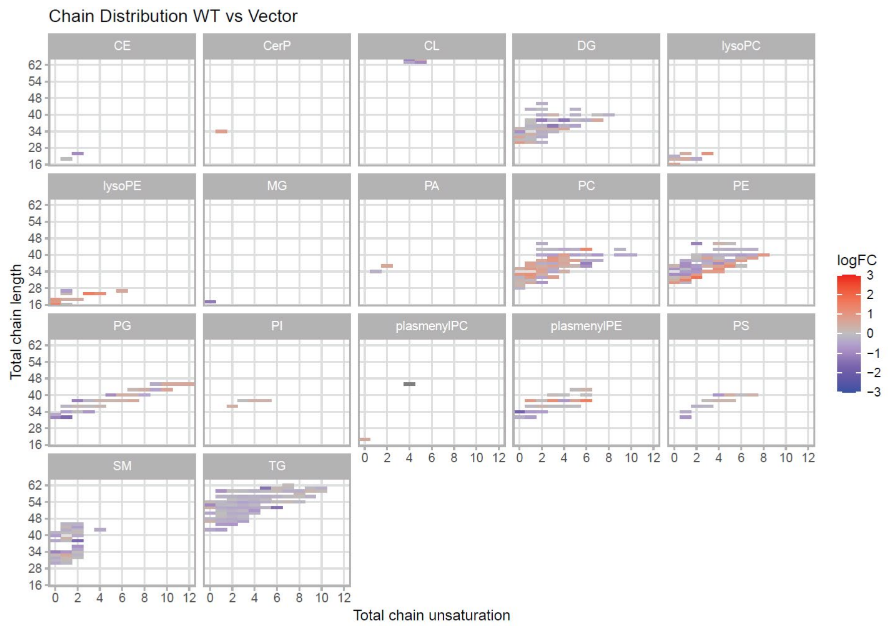


**B**

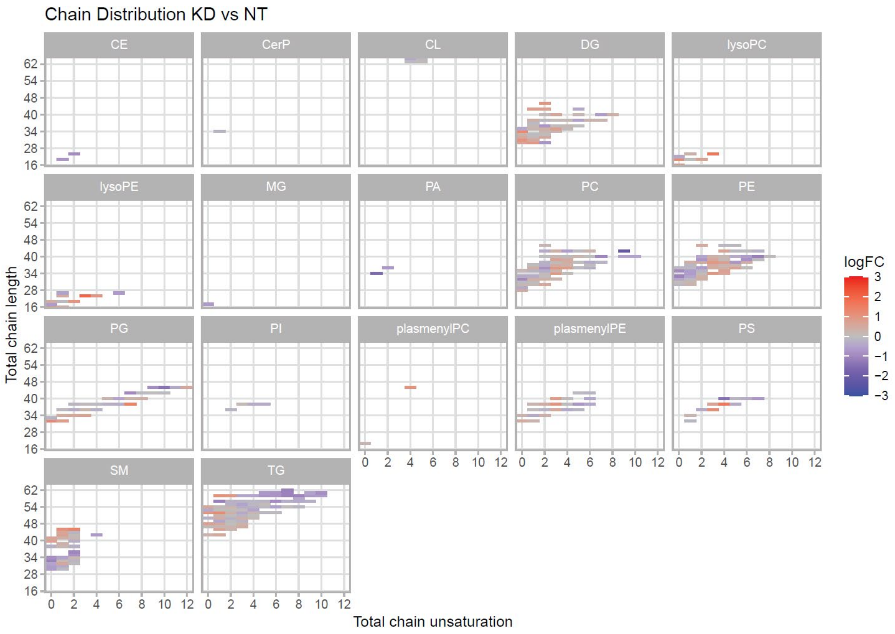


**Supplementary figure 2:** Each panel represents a lipid class and the title of the panel is the abbreviated name of the lipid class. Each box represents an individual lipid species where the colors indicate the relative ratio of the lipid (red: increased; white: no difference; blue: decreased) in (A) wild type overexpression (n=3) compared to vector control (n=3) and (B) TM6SF2 knockdown (n=5) versus non targeting control (n=4). CE (cholesterol esters), CerP (ceramide 1-phosphate), CL (Cardiolipin), DG (diacylglycerols), lysoPC (Lysophosphatidylcholines), lysoPE (lysophosphatidylethanolamine), MG (monoacylglycerols), PA (Phosphatidic acid), PC (phosphatidylcholine), PE (Phosphatidylethanolamine), PG (Phosphatidylglycerol), PI (phosphatidylinositol), plasmenylPC (Plasmenylphosphatidylcholines), plasmenylPE (Phosphatidylethanolamines), PS (Phosphatidylserines), SM (Sphingomyelines), TG (Triacylglycerols).

**Supplementary Table 1: List of lipid differentially abundant lipid species identified thorough lipidomic analysis.**

| Comparison | Molecule | Class | P.Val | Fold Change | LogFC |
|---|---|---|---|---|---|
| **KD - NT** | DG 31:0 | DG | 0.00985 | 2.09 | 1.06 |
| **KD - NT** | DG 34:0 | DG | 0.01193 | 2.35 | 1.23 |
| **KD - NT** | lysoPC 18:0 | lysoPC | 0.00007 | 2.33 | 1.22 |
| **KD - NT** | lysoPC 20:3 | lysoPC | 0.00007 | 3.03 | 1.60 |
| **KD - NT** | lysoPE 16:0 | lysoPE | 0.00433 | 2.01 | 1.00 |
| **KD - NT** | lysoPE 18:2 | lysoPE | 0.00345 | 2.01 | 1.01 |
| **KD - NT** | lysoPE 20:3 | lysoPE | 0.00007 | 3.99 | 2.00 |
| **KD - NT** | PA 34:1 | PA | 0.01258 | -2.92 | -1.55 |
| **KD - NT** | PC 42:9 | PC | 0.00262 | -4.63 | -2.21 |
| **KD - NT** | PC 40:7 | PC | 0.00013 | -2.05 | -1.03 |
| **KD - NT** | PE 40:7 | PE | 0.00011 | -2.22 | -1.15 |
| **KD - NT** | PE 33:0 | PE | 0.00099 | -2.07 | -1.05 |
| **KD - NT** | PG 44:10 | PG | 0.00024 | -2.59 | -1.38 |
| **KD - NT** | PG 32:0 | PG | 0.01088 | 2.49 | 1.31 |
| **KD - NT** | PG 38:7 | PG | 0.00007 | 2.70 | 1.43 |
| **KD - NT** | plasmenylPE 38:3 | plasmenylPE | 0.00007 | 2.04 | 1.03 |
| **KD - NT** | PS 40:4 | PS | 0.00066 | -2.69 | -1.43 |
| **KD - NT** | PS 36:3 | PS | 0.00619 | 2.28 | 1.19 |
| **KD - NT** | PS 38:4 | PS | 0.00046 | 3.12 | 1.64 |
| **KD - NT** | SM 33:0 | SM | 0.00007 | -2.10 | -1.07 |
| **KD - NT** | SM 36:2 | SM | 0.00013 | -2.04 | -1.03 |
| **KD - NT** | SM 44:1 | SM | 0.00324 | 2.11 | 1.08 |
| **KD - NT** | SM 44:2 | SM | 0.00020 | 2.60 | 1.38 |
| **KD - NT** | TG 56:1 | TG | 0.00319 | -2.23 | -1.16 |
| **KD - NT** | TG 62:7 | TG | 0.00007 | -2.18 | -1.12 |
| **KD - NT** | TG 60:7 | TG | 0.00007 | -2.14 | -1.10 |
| **KD - NT** | TG 58:7 | TG | 0.00008 | -2.13 | -1.09 |
| **KD - NT** | TG 58:8 | TG | 0.00007 | -2.08 | -1.06 |
| **KD - NT** | TG 58:1 | TG | 0.00016 | 2.13 | 1.09 |
| **KD - NT** | TG 52:1 | TG | 0.01003 | 2.30 | 1.20 |
| **WT - Vector** | CL 72:4 | CL | 0.02068 | -2.31 | -1.21 |
| **WT - Vector** | DG 38:4 | DG | 0.00272 | -2.46 | -1.30 |
| **WT - Vector** | DG 36:3 | DG | 0.00272 | -2.44 | -1.28 |
| **WT - Vector** | DG 38:2 | DG | 0.00390 | -2.35 | -1.23 |
| **WT - Vector** | lysoPE 20:4 | lysoPE | 0.00666 | 2.21 | 1.14 |
| **WT - Vector** | lysoPE 16:0 | lysoPE | 0.03878 | 2.74 | 1.46 |
| **WT - Vector** | lysoPE 20:3 | lysoPE | 0.01081 | 2.87 | 1.52 |
| **WT - Vector** | MG 17:0 | MG | 0.00390 | -2.40 | -1.26 |
| **WT - Vector** | PC 37:6 | PC | 0.02634 | -2.25 | -1.17 |
| **WT - Vector** | PC 39:3 | PC | 0.00721 | 2.04 | 1.03 |
| **WT - Vector** | PC 33:1 | PC | 0.00272 | 2.14 | 1.10 |
| **WT - Vector** | PC 42:6 | PC | 0.00882 | 2.45 | 1.29 |
| **WT - Vector** | PE 30:0 | PE | 0.00305 | 2.00 | 1.00 |

| Comparison | Molecule | Class | P.Val | Fold Change | LogFC |
|---|---|---|---|---|---|
| **WT - Vector** | PE 35:4 | PE | 0.00354 | 2.21 | 1.15 |
| **WT - Vector** | PE 31:1 | PE | 0.00199 | 2.82 | 1.50 |
| **WT - Vector** | PE 32:2 | PE | 0.00199 | 3.40 | 1.76 |
| **WT - Vector** | PG 32:1 | PG | 0.00305 | -2.96 | -1.57 |
| **WT - Vector** | PG 38:2 | PG | 0.00272 | -2.25 | -1.17 |
| **WT - Vector** | PG 32:0 | PG | 0.04293 | -2.08 | -1.06 |
| **WT - Vector** | plasmenylPC 44:4 | plasmenylPC | 0.03364 | 10.85 | 3.44 |
| **WT - Vector** | plasmenylPE 34:0 | plasmenylPE | 0.00305 | -3.46 | -1.79 |
| **WT - Vector** | plasmenylPE 38:3 | plasmenylPE | 0.00438 | 2.24 | 1.16 |
| **WT - Vector** | plasmenylPE 38:6 | plasmenylPE | 0.01815 | 2.72 | 1.44 |
| **WT - Vector** | SM 38:2 | SM | 0.03085 | -3.07 | -1.62 |
| **WT - Vector** | TG 52:6 | TG | 0.00272 | -2.71 | -1.44 |
| **WT - Vector** | TG 60:5 | TG | 0.00418 | -2.65 | -1.41 |

**Supplementary Table 1**: Only the lipid species that were differentially abundant (adjusted $p$ value < 0.05 and log fold change > 0.5 or < -0.5) in cells with either wild-type TM6SF2 overexpression or knockdown compared to their controls (vector or non-targeted controls respectively) are shown. WT (wild-type TM6SF2 overexpression, Vector (empty vector control), NT (Non-targeted shRNA control), and KD (TM6SF2 overexpression). Observed lipid species belonged to the following classes: DG (diacylglycerols), lysoPC (Lysophosphatidylcholines), lysoPE (lysophosphatidylethanolamine), MG (monoacylglycerols), PA (Phosphatidic acid), PC (phosphatidylcholine), PE (Phosphatidylethanolamine), PG (Phosphatidylglycerol), plasmenylPC (Plasmenylphosphatidylcholines), plasmenylPE (Phosphatidylethanolamines), PS (Phosphatidylserines), SM (Sphingomyelines), TG (Triacylglycerols).

**Supplementary Table 2: TM6SF2 differentially expressed lipid metabolizing genes.**

| Comparison | Symbol | Protein names | P. Val | LogFC | Fold Change |
|---|---|---|---|---|---|
| **WT - Vector** | ALDH3B1 | Aldehyde dehydrogenase family 3 member B1 | 1.00E-06 | 1.37 | 2.58 |
| **WT - Vector** | APOD | Apolipoprotein D | 1.00E-06 | 2.02 | 4.05 |
| **WT - Vector** | RGL1 | Ral guanine nucleotide dissociation stimulator-like 1 | 6.39E-06 | 0.58 | 1.49 |
| **WT - Vector** | DAB2IP | Disabled homolog 2-interacting protein | 6.54E-06 | 0.84 | 1.79 |
| **WT - Vector** | AGTR1 | Type-1 angiotensin II receptor | 2.45E-05 | -1.21 | 0.43 |
| **WT - Vector** | LIPH | Lipase member H | 7.58E-05 | 1.10 | 2.14 |
| **WT - Vector** | AKR1C1 | Aldo-keto reductase family 1 member C1 | 9.92E-05 | -1.36 | 0.39 |
| **WT - Vector** | VAV3 | Guanine nucleotide exchange factor | 0.00071 | 0.70 | 1.63 |
| **WT - Vector** | AGMO | Alkylglycerol monooxygenase | 0.00336 | -0.65 | 0.64 |
| **WT - Vector** | AKR1C4 | Aldo-keto reductase family 1 member C4 | 0.00373 | -0.98 | 0.51 |
| **WT - Vector** | FDFT1 | Squalene synthase | 0.00382 | -0.50 | 0.70 |
| **WT - Vector** | CAPN2 | Calpain-2 catalytic subunit | 0.00478 | 1.17 | 2.25 |
| **WT - Vector** | APOA2 | Apolipoprotein A-II | 0.00517 | -0.58 | 0.67 |
| **WT - Vector** | HSD17B2 | 17-beta-hydroxysteroid dehydrogenase type 2 | 0.00778 | 0.53 | 1.45 |
| **WT - Vector** | AKR1C2 | Aldo-keto reductase family 1 member C2 | 0.02397 | -1.13 | 0.46 |
| **WT - Vector** | TM6SF2 | Transmembrane 6 superfamily member 2 | 1.00E-06 | 1.75 | 3.37 |
| **KD - NT** | ACAA1 | 3-ketoacyl-CoA thiolase | 1.00E-06 | 0.81 | 1.75 |
| **KD - NT** | AKR1B1 | Aldo-keto reductase family 1 member B1 | 1.00E-06 | 2.73 | 6.64 |
| **KD - NT** | ANXA1 | Annexin A1 | 1.00E-06 | 3.79 | 13.87 |
| **KD - NT** | BMP2 | Bone morphogenetic protein 2 | 1.00E-06 | 0.94 | 1.91 |
| **KD - NT** | CYP2C19 | Cytochrome P450 2C19 | 1.00E-06 | -2.13 | 0.23 |
| **KD - NT** | EHHADH | Peroxisomal bifunctional enzyme | 1.00E-06 | -0.65 | 0.64 |
| **KD - NT** | CCN1 | CCN family member 1 | 1.00E-06 | 1.73 | 3.32 |
| **KD - NT** | MVD | Diphosphomevalonate decarboxylase | 1.00E-06 | 0.66 | 1.58 |
| **KD - NT** | PDGFRA | Platelet-derived growth factor receptor alpha | 1.00E-06 | -1.20 | 0.43 |
| **KD - NT** | PPARG | Peroxisome proliferator-activated receptor gamma | 1.00E-06 | 0.69 | 1.62 |
| **KD - NT** | PTGIS | Prostacyclin synthase | 1.00E-06 | -1.03 | 0.49 |
| **KD - NT** | THRA | Thyroid hormone receptor alpha | 1.00E-06 | -1.15 | 0.45 |
| **KD - NT** | UBE2I | SUMO-conjugating enzyme UBC9 | 1.00E-06 | 0.83 | 1.78 |
| **KD - NT** | VDR | Vitamin D3 receptor | 1.00E-06 | 1.66 | 3.16 |
| **KD - NT** | PIK3R3 | Phosphatidylinositol 3-kinase regulatory subunit gamma | 1.00E-06 | 0.89 | 1.85 |
| **KD - NT** | DGAT1 | Diacylglycerol O-acyltransferase 1 | 1.00E-06 | -0.88 | 0.55 |
| **KD - NT** | ABCC3 | ATP-binding cassette sub-family C member 3 | 1.00E-06 | 1.15 | 2.22 |
| **KD - NT** | LPCAT3 | Lysophospholipid acyltransferase 5 | 1.00E-06 | 0.54 | 1.45 |
| **KD - NT** | VAV3 | Guanine nucleotide exchange factor | 1.00E-06 | 2.22 | 4.65 |
| **KD - NT** | RGL1 | Ral guanine nucleotide dissociation stimulator-like 1 | 1.00E-06 | 0.97 | 1.96 |
| **KD - NT** | ANKRD1 | Ankyrin repeat domain-containing protein 1 | 1.00E-06 | 3.56 | 11.80 |
| **KD - NT** | TNFRSF21 | Tumor necrosis factor receptor superfamily member 21 | 1.00E-06 | 0.96 | 1.95 |
| **KD - NT** | ANGPTL3 | Angiopoietin-related protein 3 | 1.00E-06 | 1.05 | 2.08 |
| **KD - NT** | ARSJ | Arylsulfatase J | 1.00E-06 | 1.20 | 2.30 |
| **KD - NT** | ELOVL7 | Elongation of very long chain fatty acids protein 7 | 1.00E-06 | 1.26 | 2.39 |
| **KD - NT** | DGAT2 | Diacylglycerol O-acyltransferase 2 | 1.00E-06 | 0.67 | 1.59 |
| **KD - NT** | SLC44A3 | Choline transporter-like protein 3 | 1.00E-06 | 1.22 | 2.33 |
| **KD - NT** | TTC39B | Tetratricopeptide repeat protein 39B | 1.00E-06 | -1.32 | 0.40 |

| Comparison | Symbol | Protein names | P. Val | LogFC | Fold Change |
|---|---|---|---|---|---|
| **KD - NT** | AKR1C4 | Aldo-keto reductase family 1 member C4 | 1.45E-06 | 1.48 | 2.79 |
| **KD - NT** | SLCO1B3 | Solute carrier organic anion transporter family member 1B3 | 1.69E-06 | -1.35 | 0.39 |
| **KD - NT** | EPHX2 | Bifunctional epoxide hydrolase 2 | 1.91E-06 | -0.60 | 0.66 |
| **KD - NT** | SGMS1 | Phosphatidylcholine:ceramide cholinephosphotransferase 1 | 3.33E-06 | 0.82 | 1.77 |
| **KD - NT** | STUB1 | E3 ubiquitin-protein ligase | 6.40E-06 | 0.71 | 1.64 |
| **KD - NT** | ABCC1 | Multidrug resistance-associated protein 1 | 7.89E-06 | 0.87 | 1.83 |
| **KD - NT** | LIPH | Lipase member H | 7.89E-06 | 1.33 | 2.51 |
| **KD - NT** | ACBD4 | Acyl-CoA-binding domain-containing protein 4 | 1.83E-05 | -0.70 | 0.61 |
| **KD - NT** | B4GALNT1 | Beta-1,4 N-acetylgalactosaminyltransferase 1 | 2.03E-05 | 0.50 | 1.42 |
| **KD - NT** | HMGCS2 | Hydroxymethylglutaryl-CoA synthase | 2.57E-05 | 0.98 | 1.98 |
| **KD - NT** | LRP2 | Low-density lipoprotein receptor-related protein 2 | 2.84E-05 | -0.75 | 0.60 |
| **KD - NT** | GGT1 | Glutathione hydrolase 1 proenzyme | 2.85E-05 | -0.95 | 0.52 |
| **KD - NT** | PLA2G4A | Cytosolic phospholipase A2 | 2.92E-05 | -1.05 | 0.48 |
| **KD - NT** | UGT1A1 | UDP-glucuronosyltransferase 1A1 | 4.09E-05 | 1.64 | 3.13 |
| **KD - NT** | HPGD | 15-hydroxyprostaglandin dehydrogenase | 6.18E-05 | 1.18 | 2.26 |
| **KD - NT** | STARD10 | START domain-containing protein 10 | 0.000117 | -0.54 | 0.69 |
| **KD - NT** | SPTSSA | Serine palmitoyltransferase small subunit A | 0.000153 | -0.71 | 0.61 |
| **KD - NT** | PON3 | Serum paraoxonase/lactonase 3 | 0.000199 | -0.85 | 0.55 |
| **KD - NT** | B3GALNT1 | UDP-GalNAc:beta-1,3-N-acetylgalactosaminyltransferase 1 | 0.000271 | 0.61 | 1.52 |
| **KD - NT** | PPARGC1B | Peroxisome proliferator-activated receptor gamma coactivator 1-beta | 0.000291 | -0.61 | 0.66 |
| **KD - NT** | LPCAT4 | Lysophospholipid acyltransferase LPCAT4 | 0.000433 | -0.77 | 0.59 |
| **KD - NT** | GDPD1 | Lysophospholipase D GDPD1 | 0.000553 | -0.55 | 0.68 |
| **KD - NT** | SLC27A2 | Very long-chain acyl-CoA synthetase | 0.000665 | -0.85 | 0.55 |
| **KD - NT** | OSBPL1A | Oxysterol-binding protein-related protein 1 | 0.000748 | -0.74 | 0.60 |
| **KD - NT** | CYP2R1 | Vitamin D 25-hydroxylase | 0.000748 | -0.58 | 0.67 |
| **KD - NT** | DAB2IP | Disabled homolog 2-interacting protein | 0.000832 | -0.65 | 0.64 |
| **KD - NT** | CAPN2 | Calpain-2 catalytic subunit | 0.000931 | 1.31 | 2.48 |
| **KD - NT** | ACP6 | Lysophosphatidic acid phosphatase type 6 | 0.000962 | -0.53 | 0.69 |
| **KD - NT** | SEC14L2 | SEC14-like protein 2 | 0.001312 | -0.84 | 0.56 |
| **KD - NT** | APOA5 | Apolipoprotein A-V | 0.001321 | -1.12 | 0.46 |
| **KD - NT** | TGFB1 | Transforming growth factor beta-1 proprotein | 0.001356 | -0.90 | 0.54 |
| **KD - NT** | LSR | Lipolysis-stimulated lipoprotein receptor | 0.001498 | 0.51 | 1.43 |
| **KD - NT** | PLIN5 | Perilipin-5 | 0.001606 | -0.90 | 0.54 |
| **KD - NT** | SLCO1A2 | Solute carrier organic anion transporter family member 1A2 | 0.001779 | -0.63 | 0.65 |
| **KD - NT** | G0S2 | G0/G1 switch protein 2 | 0.002284 | -0.91 | 0.53 |
| **KD - NT** | SPHK1 | Sphingosine kinase 1 | 0.002309 | -0.53 | 0.69 |
| **KD - NT** | SIRT3 | NAD-dependent protein deacetylase sirtuin-3 | 0.002604 | -0.51 | 0.70 |
| **KD - NT** | SULT2A1 | Sulfotransferase 2A1 | 0.002642 | -0.60 | 0.66 |
| **KD - NT** | DECR2 | Peroxisomal 2,4-dienoyl-CoA reductase | 0.002888 | 0.56 | 1.48 |
| **KD - NT** | PON1 | Serum paraoxonase/arylesterase 1 | 0.003019 | -0.77 | 0.59 |
| **KD - NT** | PNPLA2 | Patatin-like phospholipase domain-containing protein 2 | 0.003203 | -0.69 | 0.62 |
| **KD - NT** | PLA2G6 | Patatin-like phospholipase domain-containing protein 9 | 0.003506 | -0.85 | 0.55 |
| **KD - NT** | ACOT7 | Cytosolic acyl coenzyme A thioester hydrolase | 0.003515 | 0.53 | 1.44 |
| **KD - NT** | EGR1 | Early growth response protein 1 | 0.003783 | 0.84 | 1.78 |
| **KD - NT** | PDK1 | 3-phosphoinositide-dependent protein kinase 1 | 0.004379 | -0.63 | 0.64 |
| **KD - NT** | PDK1 | Pyruvate dehydrogenase kinase isoform 1 | 0.004379 | -0.63 | 0.64 |
| **KD - NT** | SLC45A3 | Solute carrier family 45 member 3 | 0.004518 | 0.88 | 1.85 |

| Comparison | Symbol | Protein names | P. Val | LogFC | Fold Change |
|---|---|---|---|---|---|
| **KD - NT** | PDGFB | Platelet-derived growth factor subunit B | 0.005481 | 1.29 | 2.45 |
| **KD - NT** | ID2 | DNA-binding protein inhibitor ID-2 | 0.005720 | 0.77 | 1.71 |
| **KD - NT** | MED4 | Mediator complex subunit 4 | 0.006551 | -0.58 | 0.67 |
| **KD - NT** | INPP5J | Inositol polyphosphate 5-phosphatase J | 0.008313 | 1.12 | 2.17 |
| **KD - NT** | AKR1D1 | Aldo-keto reductase family 1 member D1 | 0.008674 | 1.33 | 2.52 |
| **KD - NT** | PDGFA | Platelet-derived growth factor subunit A | 0.008825 | -0.55 | 0.68 |
| **KD - NT** | APOC3 | Apolipoprotein C-III | 0.009457 | -0.87 | 0.55 |
| **KD - NT** | GNB3 | Guanine nucleotide-binding protein G | 0.010164 | -0.85 | 0.55 |
| **KD - NT** | ALB | Albumin | 0.010572 | 0.55 | 1.46 |
| **KD - NT** | ALB | Fas-binding factor 1 | 0.010572 | 0.55 | 1.46 |
| **KD - NT** | MED7 | Mediator of RNA polymerase II transcription subunit 7 | 0.011572 | -0.54 | 0.69 |
| **KD - NT** | HSD17B8 | Estradiol 17-beta-dehydrogenase 8 | 0.013272 | -0.71 | 0.61 |
| **KD - NT** | LDLR | Low-density lipoprotein receptor | 0.015460 | 0.55 | 1.47 |
| **KD - NT** | ABHD4 | Alpha/beta hydrolase domain-containing protein 4 | 0.017082 | 0.56 | 1.47 |
| **KD - NT** | FDX1 | Ferredoxin-1 | 0.020050 | 0.56 | 1.48 |
| **KD - NT** | CIDEC | Cell death activator CIDE-3 | 0.028195 | 0.87 | 1.83 |
| **KD - NT** | CYP2C9 | Cytochrome P450 2C9 | 0.029385 | -0.99 | 0.50 |
| **KD - NT** | ACER3 | Alkaline ceramidase 3 | 0.031183 | 0.52 | 1.43 |
| **KD - NT** | AKR1C2 | Aldo-keto reductase family 1 member C2 | 0.037809 | 1.00 | 2.00 |
| **KD - NT** | TM6SF2 | Transmembrane 6 superfamily member 2 | 0.428827 | -0.29 | 0.82 |

**Supplementary Table 2:** Genes that were differentially expressed **(**adjusted *p* value of less than 0.05 and an absolute log fold change of greater than 1 or 1.4-fold change) in cells with either wild-type TM6SF2 overexpression or knockdown compared to their controls (vector or non-targeted controls respectively) are shown. Only the genes that WT (wild-type TM6SF2 overexpression, Vector (empty vector control), NT (Non-targeted shRNA control), and KD (TM6SF2 overexpression). Only the set of 1008 genes for the 749 proteins identified by Reactome as being part of the overall human Metabolism of lipids pathway (R-HSA-556833) and the 864 human proteins identified in GO as being involved in the regulation of lipid metabolic process (GO:0019216) were analyzed.
