## Supplemental cell profiler method for "Perturbation of transmembrane 6 superfamily member 2 expression alters lipid metabolism in a human liver cell line"

CellProfiler Pipeline: http://www.cellprofiler.org
Version:5
DateRevision:413
GitHash:
ModuleCount:22
HasImagePlaneDetails:False

Images:[module_num:1|svn_version:'Unknown'|variable_revision_number:2|show_window:False|notes:['To begin creating your project, use the Images module to compile a list of files and/or folders that you want to analyze. You can also specify a set of rules to include only the desired files in your selected folders.']|batch_state:array([], dtype=uint8)|enabled:True|wants_pause:False]
    :
    Filter images?:Custom
    Select the rule criteria:and (file does endwith ".C01")

Metadata:[module_num:2|svn_version:'Unknown'|variable_revision_number:6|show_window:False|notes:['The Metadata module optionally allows you to extract information describing your images (i.e, metadata) which will be stored along with your measurements. This information can be contained in the file name and/or location, or in an external file.']|batch_state:array([], dtype=uint8)|enabled:True|wants_pause:False]
    Extract metadata?:Yes
    Metadata data type:Text
    Metadata types:{}
    Extraction method count:1
    Metadata extraction method:Extract from file/folder names
    Metadata source:File name
    Regular expression to extract from file name:^(?P<WELL>.*)_(?P<CELL>.*)_(?P<COND>.*)_(?P<srs>.*)_(?P<dye>.*).C01
    Regular expression to extract from folder name:(?P<Date>[0-9]{4}_[0-9]{2}_[0-9]{2})$
    Extract metadata from:All images
    Select the filtering criteria:and (file does contain "")
    Metadata file location:Elsewhere...|
    Match file and image metadata:[]
    Use case insensitive matching?:No
    Metadata file name:
    Does cached metadata exist?:No

NamesAndTypes:[module_num:3|svn_version:'Unknown'|variable_revision_number:8|show_window:False|notes:['The NamesAndTypes module allows you to assign a meaningful name to each image by which other modules will refer to it.']|batch_state:array([], dtype=uint8)|enabled:True|wants_pause:False]
    Assign a name to:Images matching rules
    Select the image type:Grayscale image
    Name to assign these images:DNA
    Match metadata:[{'Raw_Lipid': 'CELL', 'Raw_DNA': 'CELL'}, {'Raw_Lipid': 'COND', 'Raw_DNA': 'COND'}, {'Raw_Lipid': 'WELL', 'Raw_DNA': 'WELL'}, {'Raw_Lipid': 'srs', 'Raw_DNA': 'srs'}]
    Image set matching method:Metadata
    Set intensity range from:Image metadata
    Assignments count:2
    Single images count:0
    Maximum intensity:255.0
    Process as 3D?:No
    Relative pixel spacing in X:1.0
    Relative pixel spacing in Y:1.0
    Relative pixel spacing in Z:1.0
    Select the rule criteria:and (file does endwith "DNA.C01")
    Name to assign these images:Raw_DNA
    Name to assign these objects:Cell
    Select the image type:Grayscale image
    Set intensity range from:Image metadata
    Maximum intensity:255.0
    Select the rule criteria:and (file does endwith "Lipid.C01")
    Name to assign these images:Raw_Lipid
    Name to assign these objects:Nucleus
    Select the image type:Grayscale image
    Set intensity range from:Image metadata
    Maximum intensity:255.0

Groups:[module_num:4|svn_version:'Unknown'|variable_revision_number:2|show_window:False|notes:['The Groups module optionally allows you to split your list of images into image subsets (groups) which will be processed independently of each other. Examples of groupings include screening batches, microtiter plates, time-lapse movies, etc.']|batch_state:array([], dtype=uint8)|enabled:True|wants_pause:False]
    Do you want to group your images?:No
    grouping metadata count:1
    Metadata category:None

RescaleIntensity:[module_num:5|svn_version:'Unknown'|variable_revision_number:3|show_window:False|notes:[]|batch_state:array([], dtype=uint8)|enabled:True|wants_pause:False]
    Select the input image:Raw_DNA
    Name the output image:DNA
    Rescaling method:Stretch each image to use the full intensity range
    Method to calculate the minimum intensity:Custom
    Method to calculate the maximum intensity:Custom
    Lower intensity limit for the input image:0.0
    Upper intensity limit for the input image:1.0
    Intensity range for the input image:0.0,1.0
    Intensity range for the output image:0.0,1.0
    Select image to match in maximum intensity:None
    Divisor value:1.0
    Divisor measurement:None

RescaleIntensity:[module_num:6|svn_version:'Unknown'|variable_revision_number:3|show_window:False|notes:[]|batch_state:array([], dtype=uint8)|enabled:True|wants_pause:False]
    Select the input image:Raw_Lipid
    Name the output image:Lipid
    Rescaling method:Match the image's maximum to another image's maximum
    Method to calculate the minimum intensity:Custom
    Method to calculate the maximum intensity:Custom
    Lower intensity limit for the input image:0.0
    Upper intensity limit for the input image:1.0
    Intensity range for the input image:0.0,1.0
    Intensity range for the output image:0.0,1.0
    Select image to match in maximum intensity:DNA
    Divisor value:1.0
    Divisor measurement:None

EnhanceOrSuppressFeatures:[module_num:7|svn_version:'Unknown'|variable_revision_number:7|show_window:False|notes:[]|batch_state:array([], dtype=uint8)|enabled:True|wants_pause:False]
    Select the input image:Lipid
    Name the output image:Enhanced_Lipid
    Select the operation:Enhance
    Feature size:30
    Feature type:Speckles
    Range of hole sizes:1,10
    Smoothing scale:2.0
    Shear angle:0.0
    Decay:0.95
    Enhancement method:Tubeness
    Speed and accuracy:Fast
    Rescale result image:Yes

IdentifyPrimaryObjects:[module_num:8|svn_version:'Unknown'|variable_revision_number:14|show_window:False|notes:[]|batch_state:array([], dtype=uint8)|enabled:True|wants_pause:False]
    Select the input image:DNA
    Name the primary objects to be identified:Nuclei
    Typical diameter of objects, in pixel units (Min,Max):50,250
    Discard objects outside the diameter range?:Yes
    Discard objects touching the border of the image?:Yes
    Method to distinguish clumped objects:Intensity
    Method to draw dividing lines between clumped objects:Intensity
    Size of smoothing filter:10
    Suppress local maxima that are closer than this minimum allowed distance:7.0
    Speed up by using lower-resolution image to find local maxima?:Yes
    Fill holes in identified objects?:After both thresholding and declumping
    Automatically calculate size of smoothing filter for declumping?:Yes
    Automatically calculate minimum allowed distance between local maxima?:Yes
    Handling of objects if excessive number of objects identified:Continue
    Maximum number of objects:500
    Display accepted local maxima?:No
    Select maxima color:Blue
    Use advanced settings?:Yes
    Threshold setting version:12
    Threshold strategy:Global
    Thresholding method:Minimum Cross-Entropy
    Threshold smoothing scale:2
    Threshold correction factor:2
    Lower and upper bounds on threshold:0.17,1
    Manual threshold:0.0
    Select the measurement to threshold with:None
    Two-class or three-class thresholding?:Two classes
    Log transform before thresholding?:No
    Assign pixels in the middle intensity class to the foreground or the background?:Foreground
    Size of adaptive window:50
    Lower outlier fraction:0.05
    Upper outlier fraction:0.05
    Averaging method:Mean
    Variance method:Standard deviation
    # of deviations:2.0
    Thresholding method:Otsu

IdentifyPrimaryObjects:[module_num:9|svn_version:'Unknown'|variable_revision_number:14|show_window:False|notes:[]|batch_state:array([], dtype=uint8)|enabled:True|wants_pause:False]
    Select the input image:Enhanced_Lipid
    Name the primary objects to be identified:Lipid_Droplets
    Typical diameter of objects, in pixel units (Min,Max):1,30
    Discard objects outside the diameter range?:Yes
    Discard objects touching the border of the image?:Yes
    Method to distinguish clumped objects:Intensity
    Method to draw dividing lines between clumped objects:Propagate
    Size of smoothing filter:10
    Suppress local maxima that are closer than this minimum allowed distance:7.0
    Speed up by using lower-resolution image to find local maxima?:Yes
    Fill holes in identified objects?:After both thresholding and declumping
    Automatically calculate size of smoothing filter for declumping?:Yes
    Automatically calculate minimum allowed distance between local maxima?:Yes
    Handling of objects if excessive number of objects identified:Continue
    Maximum number of objects:500
    Display accepted local maxima?:No
    Select maxima color:Blue
    Use advanced settings?:Yes
    Threshold setting version:12
    Threshold strategy:Global
    Thresholding method:Minimum Cross-Entropy
    Threshold smoothing scale:1
    Threshold correction factor:2
    Lower and upper bounds on threshold:0.05,0.5
    Manual threshold:0.0
    Select the measurement to threshold with:None
    Two-class or three-class thresholding?:Two classes
    Log transform before thresholding?:No
    Assign pixels in the middle intensity class to the foreground or the background?:Foreground
    Size of adaptive window:50
    Lower outlier fraction:0.05
    Upper outlier fraction:0.05
    Averaging method:Mean
    Variance method:Standard deviation
    # of deviations:2.0
    Thresholding method:Otsu

IdentifySecondaryObjects:[module_num:10|svn_version:'Unknown'|variable_revision_number:10|show_window:False|notes:[]|batch_state:array([], dtype=uint8)|enabled:True|wants_pause:False]
    Select the input objects:Nuclei
    Name the objects to be identified:Lipid_Cells
    Select the method to identify the secondary objects:Propagation
    Select the input image:Lipid
    Number of pixels by which to expand the primary objects:10
    Regularization factor:0.05
    Discard secondary objects touching the border of the image?:Yes
    Discard the associated primary objects?:Yes
    Name the new primary objects:LipidCellNoNuclei
    Fill holes in identified objects?:No
    Threshold setting version:12
    Threshold strategy:Global
    Thresholding method:Minimum Cross-Entropy
    Threshold smoothing scale:0.5
    Threshold correction factor:.9
    Lower and upper bounds on threshold:0.06,1.0
    Manual threshold:0.0
    Select the measurement to threshold with:None
    Two-class or three-class thresholding?:Two classes
    Log transform before thresholding?:No
    Assign pixels in the middle intensity class to the foreground or the background?:Foreground
    Size of adaptive window:50
    Lower outlier fraction:0.05
    Upper outlier fraction:0.05
    Averaging method:Mean
    Variance method:Standard deviation
    # of deviations:2.0
    Thresholding method:Otsu

IdentifyTertiaryObjects:[module_num:11|svn_version:'Unknown'|variable_revision_number:3|show_window:False|notes:[]|batch_state:array([], dtype=uint8)|enabled:True|wants_pause:False]
    Select the larger identified objects:Lipid_Cells
    Select the smaller identified objects:Nuclei
    Name the tertiary objects to be identified:perimeter
    Shrink smaller object prior to subtraction?:Yes

RelateObjects:[module_num:12|svn_version:'Unknown'|variable_revision_number:5|show_window:False|notes:[]|batch_state:array([], dtype=uint8)|enabled:True|wants_pause:False]
    Parent objects:Lipid_Cells
    Child objects:Lipid_Droplets
    Calculate child-parent distances?:None
    Calculate per-parent means for all child measurements?:Yes
    Calculate distances to other parents?:No
    Do you want to save the children with parents as a new object set?:Yes
    Name the output object:RelateObjects_lipidCellLipidDroplets
    Parent name:None
    Parent name:None
    Parent name:None
    Parent name:None
    Parent name:None
    Parent name:None
    Parent name:None
    Parent name:None
    Parent name:None
    Parent name:None
    Parent name:None
    Parent name:None
    Parent name:None
    Parent name:None
    Parent name:None
    Parent name:None
    Parent name:None
    Parent name:None
    Parent name:None
    Parent name:None
    Parent name:None
    Parent name:None
    Parent name:None
    Parent name:None
    Parent name:None
    Parent name:None
    Parent name:None
    Parent name:None
    Parent name:None

MeasureObjectIntensity:[module_num:13|svn_version:'Unknown'|variable_revision_number:4|show_window:False|notes:[]|batch_state:array([], dtype=uint8)|enabled:True|wants_pause:False]
    Select images to measure:Raw_Lipid
    Select objects to measure:Lipid_Cells, Lipid_Droplets

MeasureObjectIntensity:[module_num:14|svn_version:'Unknown'|variable_revision_number:4|show_window:False|notes:[]|batch_state:array([], dtype=uint8)|enabled:True|wants_pause:False]
    Select images to measure:Lipid
    Select objects to measure:Lipid_Cells, Lipid_Droplets

MeasureImageIntensity:[module_num:15|svn_version:'Unknown'|variable_revision_number:4|show_window:False|notes:[]|batch_state:array([], dtype=uint8)|enabled:True|wants_pause:False]
    Select images to measure:Lipid
    Measure the intensity only from areas enclosed by objects?:Yes
    Select input object sets:Lipid_Cells
    Calculate custom percentiles:No
    Specify percentiles to measure:10,90

CalculateMath:[module_num:16|svn_version:'Unknown'|variable_revision_number:3|show_window:False|notes:[]|batch_state:array([], dtype=uint8)|enabled:True|wants_pause:False]
    Name the output measurement:Total_Frame_Lipid_Intensity_per_Cell
    Operation:Divide
    Select the numerator measurement type:Image
    Select the numerator objects:None
    Select the numerator measurement:Intensity_MeanIntensity_Lipid_Lipid_Cells
    Multiply the above operand by:1.0
    Raise the power of above operand by:1.0
    Select the denominator measurement type:Object
    Select the denominator objects:Nuclei
    Select the denominator measurement:Number_Object_Number
    Multiply the above operand by:1.0
    Raise the power of above operand by:1.0
    Take log10 of result?:No
    Multiply the result by:1.0
    Raise the power of result by:1.0
    Add to the result:0.0
    How should the output value be rounded?:Not rounded
    Enter how many decimal places the value should be rounded to:0
    Constrain the result to a lower bound?:No
    Enter the lower bound:0.0
    Constrain the result to an upper bound?:No
    Enter the upper bound:1.0

GrayToColor:[module_num:17|svn_version:'Unknown'|variable_revision_number:4|show_window:False|notes:[]|batch_state:array([], dtype=uint8)|enabled:True|wants_pause:False]
    Select a color scheme:RGB
    Rescale intensity:No
    Select the image to be colored red:Leave this black
    Select the image to be colored green:Lipid
    Select the image to be colored blue:DNA
    Name the output image:MergedColorImage
    Relative weight for the red image:1.0
    Relative weight for the green image:1.0
    Relative weight for the blue image:1.0
    Select the image to be colored cyan:Leave this black
    Select the image to be colored magenta:Leave this black
    Select the image to be colored yellow:Leave this black
    Select the image that determines brightness:Leave this black
    Relative weight for the cyan image:1.0
    Relative weight for the magenta image:1.0
    Relative weight for the yellow image:1.0
    Relative weight for the brightness image:1.0
    Hidden:1
    Image name:None
    Color:#ff0000
    Weight:1.0

OverlayOutlines:[module_num:18|svn_version:'Unknown'|variable_revision_number:4|show_window:False|notes:[]|batch_state:array([], dtype=uint8)|enabled:True|wants_pause:False]
    Display outlines on a blank image?:Yes
    Select image on which to display outlines:None
    Name the output image:ObjectOutlines
    Outline display mode:Color
    Select method to determine brightness of outlines:Max of image
    How to outline:Thick
    Select outline color:#0080FF
    Select objects to display:Nuclei
    Select outline color:yellow
    Select objects to display:Lipid_Cells
    Select outline color:green
    Select objects to display:Lipid_Droplets

Tile:[module_num:19|svn_version:'Unknown'|variable_revision_number:1|show_window:True|notes:[]|batch_state:array([], dtype=uint8)|enabled:True|wants_pause:False]
    Select an input image:DNA
    Name the output image:TiledResults
    Tile assembly method:Within cycles
    Final number of rows:1
    Final number of columns:3
    Image corner to begin tiling:top left
    Direction to begin tiling:row
    Use meander mode?:No
    Automatically calculate number of rows?:No
    Automatically calculate number of columns?:No
    Select an additional image to tile:Enhanced_Lipid
    Select an additional image to tile:ObjectOutlines

RescaleIntensity:[module_num:20|svn_version:'Unknown'|variable_revision_number:3|show_window:False|notes:[]|batch_state:array([], dtype=uint8)|enabled:True|wants_pause:False]
    Select the input image:TiledResults
    Name the output image:RescaledTiledResults
    Rescaling method:Match the image's maximum to another image's maximum
    Method to calculate the minimum intensity:Custom
    Method to calculate the maximum intensity:Custom
    Lower intensity limit for the input image:0.0
    Upper intensity limit for the input image:1.0
    Intensity range for the input image:0.0,1.0
    Intensity range for the output image:0.0,1.0
    Select image to match in maximum intensity:DNA
    Divisor value:1.0
    Divisor measurement:None

SaveImages:[module_num:21|svn_version:'Unknown'|variable_revision_number:15|show_window:False|notes:[]|batch_state:array([], dtype=uint8)|enabled:False|wants_pause:False]
    Select the type of image to save:Image
    Select the image to save:RescaledTiledResults
    Select method for constructing file names:Single name
    Select image name for file prefix:None
    Enter single file name:\g<CELL>_\g<COND>_\g<WELL>_\g<srs>
    Number of digits:4
    Append a suffix to the image file name?:No
    Text to append to the image name:
    Saved file format:tiff
    Output file location:Elsewhere...|E:\\Speliotes\\Lipid_Staining_Pipeline\\TM6SF2\\2020_02_06_TM6SF2_KD_Nof1\\CellProfilerResultsRenamedCorrectlyRun1
    Image bit depth:8-bit integer
    Overwrite existing files without warning?:Yes
    When to save:Every cycle
    Record the file and path information to the saved image?:Yes
    Create subfolders in the output folder?:No
    Base image folder:Elsewhere...|
    How to save the series:T (Time)

ExportToSpreadsheet:[module_num:22|svn_version:'Unknown'|variable_revision_number:13|show_window:False|notes:[]|batch_state:array([], dtype=uint8)|enabled:True|wants_pause:False]
    Select the column delimiter:Comma (",")
    Add image metadata columns to your object data file?:Yes
    Add image file and folder names to your object data file?:No
    Select the measurements to export:No
    Calculate the per-image mean values for object measurements?:Yes
    Calculate the per-image median values for object measurements?:No
    Calculate the per-image standard deviation values for object measurements?:No
    Output file location:Elsewhere...|E:\\Speliotes\\Lipid_Staining_Pipeline\\TM6SF2\\2020_02_06_TM6SF2_KD_Nof1\\CellProfilerResultsRenamedCorrectlyRun1\\Excel
    Create a GenePattern GCT file?:No
    Select source of sample row name:Metadata
    Select the image to use as the identifier:None
    Select the metadata to use as the identifier:None
    Export all measurement types?:Yes
    Press button to select measurements:
    Representation of Nan/Inf:NaN
    Add a prefix to file names?:Yes
    Filename prefix:V3_
    Overwrite existing files without warning?:Yes
    Data to export:Do not use
    Combine these object measurements with those of the previous object?:No
    File name:DATA.csv
    Use the object name for the file name?:Yes
